# Characterization of eight new *Hydractinia* i-cell markers reveals underlying heterogeneity in the adult pluripotent stem cell population

**DOI:** 10.1101/2024.07.07.602406

**Authors:** Justin Waletich, Danielle de Jong, Christine E. Schnitzler

## Abstract

Adult pluripotent stem cells are found in diverse animals, including cnidarians, acoels, and planarians, and confer remarkable abilities such as whole-body regeneration. The mechanisms by which these pluripotent stem cells orchestrate the replacement of all lost cell types, however, remains poorly understood. Underlying heterogeneity within the stem cell populations of these animals is often obscured when focusing on certain tissue types or life history stages, which tend to have indistinguishable spatial expression patterns of stem cell marker genes. Here, we focus on the adult pluripotent stem cells (i-cells) of *Hydractinia symbiolongicarpus*, a colonial marine cnidarian with distinct polyp types and stolonal tissue. Recently, a single-cell expression atlas was generated for *H. symbiolongicarpus* which revealed two distinct clusters with i-cell signatures, potentially representing heterogeneity within this species’ stem cell population. Considering this finding, we investigated eight new putative stem cell marker genes from the atlas including five expressed in both i-cell clusters (*Pcna*, *Nop58*, *Mcm4*, *Ubr7*, and *Uhrf1*) and three expressed in one cluster or the other (*Pter, FoxQ2-like,* and *Zcwpw1*). We characterized their expression patterns in various contexts *–* feeding and sexual polyps, juvenile feeding polyps, stolon, and during feeding polyp head regeneration *–* revealing context-dependent gene expression patterns and a transcriptionally dynamic i-cell population. We uncover previously unknown differences within the i-cell population of *Hydractinia* and demonstrate that its colonial nature serves as an excellent system for investigating and visualizing heterogeneity in pluripotent stem cells.

## Introduction

Pluripotent stem cells are self-renewing cells, capable of differentiating into all cell types in an organism [1]. Unlike in mammals, where these cells are limited to embryonic development, some invertebrates maintain a population of pluripotent stem cells into adulthood, enabling the indefinite maintenance and regeneration of complex structures or whole animals [2–5]. Well-known examples include neoblasts in planarians and acoels, and i-cells in cnidarians [6–8]. These adult pluripotent stem cells express high levels of *Piwi*, a gene originally characterized as a prominent member of the germline multipotency program [2,5,9–11]. *Piwi* expression is not restricted to germline cells or stem cells, however, as it has been shown to be broadly expressed in some somatic tissues across several animals [12–14]. Additionally, recent work suggests the stem cells of planarians, acoels, and annelids are transcriptionally dynamic and show *Piwi* expression in progenitor and some non-cycling cell types [3,5,15,16]. Modern techniques like single-cell transcriptomics are generating a wealth of new data, including new genetic markers of stem cell populations beyond the well-known stem cell markers [17–22]. Recent studies characterizing stem cell clusters and their genetic markers in animals with adult pluripotent and multipotent stem cells are providing valuable insights into the remarkable ability of these cells to become all cell types within the organism [3,5,15,19,21].

The phylum Cnidaria consists of a diverse group of aquatic invertebrates including corals, sea anemones, jellyfish, and hydroids, many of which have been shown to possess adult stem cells [23–26]. Hydrozoan i-cells are the most studied among the adult stem cells in cnidarians, with their name referring to their location in the interstitial spaces of the ectodermal epithelium [27]. *Hydractinia* is currently the only hydrozoan with i-cells that have been shown to be pluripotent [8]. As a colonial marine invertebrate, it has several morphologically distinct and functionally specialized polyp types that are connected through a basal mat tissue called stolon (Fig 1A) [28]. *Hydractinia* harbors i-cells in multiple sites within the colony that can be identified based on a combination of characteristics such as high *Piwi1* expression, a large nuclear-to-cytoplasmic ratio, 7-10µm cell diameter, and the presence of large amounts of ribosomes [7,27,29]. The stolon contains a large population of i-cells that are ubiquitously dispersed, while *Hydractinia* feeding polyps have a characteristic population of proliferating i-cells in a band region in the lower body column (Fig 1B) [2,29]. In stolon and feeding polyps, areas adjacent to i-cells are the sites of both neurogenesis and nematogenesis [29–31]. I-cells are also present in sexual polyps, where a subset migrate from the ectodermal layer to the endodermal layer in an area known as the germinal zone, and eventually become gametes in specialized compartments of the sporosacs (Fig 1C) [2,25,32–35].

**Fig 1.**
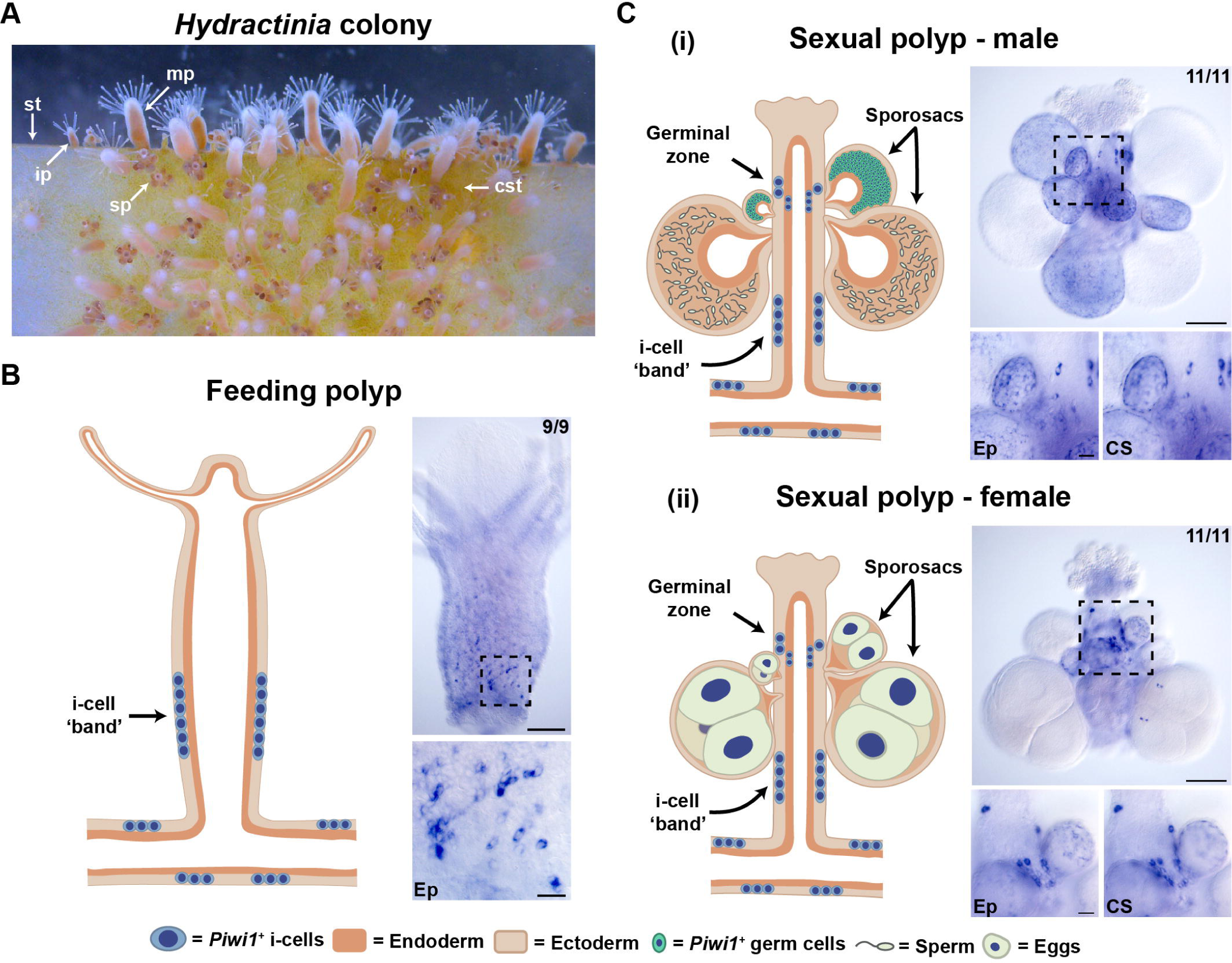
*Piwi1* spatial expression patterns in *Hydractinia* feeding and sexual polyps showing i-cell locations in the colony. (A) Image of a *Hydractinia* colony growing on a glass slide. mp: mature feeding polyp, ip: immature feeding polyp, sp: sexual polyp, st; stolon, cst: chitinous stolon. (B) Schematic (left) and colorimetric *in situ* hybridization images (right) showing i-cell locations in *Hydractinia* feeding polyps and stolon as seen with *Piwi1* expression. (C) Same as in (B) but with male (i) and female (ii) sexual polyps and stolon. For each polyp type, the black dashed box in the upper *in situ* image indicates the region selected for higher magnification in the lower *in situ* images. Upper *in situ* images have 100µm scale bars and lower *in situ* images have 20µm scale bars. Numbers in the top right corner show the proportion of samples that reflect the image shown. Ep is the Ectodermal plane; CS is the cross-section plane.

*Hydractinia* possesses incredible regenerative abilities that have been studied for well over a century, and i-cells and progenitors are critical for these capabilities [29,36–39]. Decapitated feeding polyps undergo three main phases of oral regeneration, including wound closure over the injury within 4 hours post-decapitation (hpd), establishment of a blastema in the region of the wound around 24hpd, and finally formation of a fully functional head with tentacles capable of prey capture between 48-72hpd [29,31,39–41]. *Piwi1^+^* i-cells migrate from the lower body column to the oral end to contribute to the formation of the blastema [29].

Recently, single-cell transcriptomics has been used to identify distinct cell types and cell states from dissociated whole animals or tissues, based on differentially expressed genetic markers [42]. The published *Hydractinia* single-cell expression atlas included in the genome paper [43] includes 8,800 cells forming 18 different cell types, including two separate i-cell/progenitor cell clusters: one fated towards somatic lineages, and one fated towards germ cell lineages (Fig 2A). These clusters were annotated as i-cells/progenitor cells based on the expression of known i-cell genes such as *Piwi1*, *Piwi2*, *Polynem*, *Vasa, Pl10, SoxB1*, and *GNL3*, though these genes have somewhat ‘leaky’ expression in the cellular atlas and are also present in many other cell type clusters (Fig 2C and S1 Fig) [43]. The expression of known stem cell markers like *Piwi1* in non-stem cell clusters is also found in the *Hydra* and *Clytia* single-cell atlases, as well as in atlases from other non-cnidarian invertebrate models with adult pluripotent stem cells, such as planarians and acoels [3,15,22,44]. One hypothesis is that *Piwi^+^* stem cells represent a mixed population of distinct stem cell subtypes, including progenitor populations. Whether the two i-cell clusters in the *Hydractinia* single-cell atlas represent distinct i-cell subtypes has yet to be thoroughly characterized and is one of the open questions we sought to investigate.

**Fig 2.**
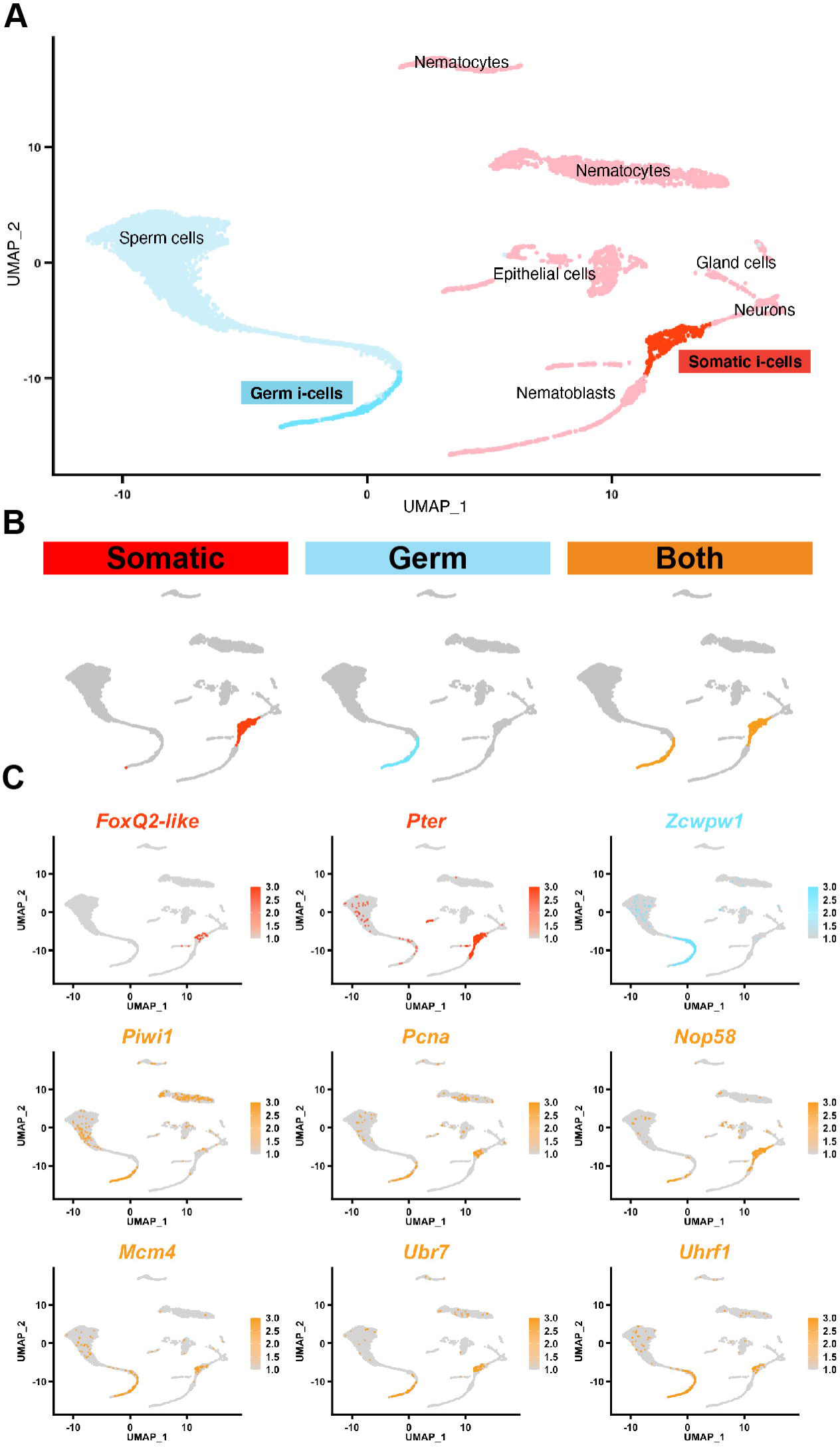
New marker genes identified from the two i-cell clusters in the *Hydractinia* single-cell atlas. (A) *Hydractinia* single-cell atlas with simplified cluster names, colored by somatic (pink and red) and germ (blue) lineages [43]. The two i-cell clusters are labeled ‘Somatic i-cells’ and ‘Germ i-cells’. (B) The strategy to identify new i-cell markers from the atlas is shown by highlighting the ‘Somatic i-cells’ cluster in red, the ‘Germ i-cells’ cluster in blue, and both i-cell clusters in gold. (C) Expression maps for targeted i-cell genes in the *Hydractinia* single-cell atlas. Each gene is highlighted by the color (red, blue, or gold from B) that represents the strategy used to identify it.

Here, we further explore the *Hydractinia symbiolongicarpus* single-cell atlas by selecting a set of genetic markers that are expressed in the two i-cell clusters. By taking advantage of the unique biology of *Hydractinia –* the ability to study different polyp types, different developmental stages of feeding polyps, stolon, and feeding polyp head regeneration in a single system *–* we acquire a better understanding of the possible roles of the new markers in the colony and depict an adult stem cell population that is characterized by context-dependent gene expression patterns. We show that our new marker genes and *Piwi1* have consistent patterns of expression in the most commonly studied areas of the colony known to possess i-cells, such as in the i-cell band of adult feeding polyps and in stolonal tissue, concealing meaningful differences within the i-cell population across the colony. This contrasts with the results we obtained when examining different biological contexts, such as in sexual polyps, in different stages of feeding polyp development, and during feeding polyp head regeneration, where we uncovered variable expression patterns of these genes. These results suggest that the two i-cell clusters in the atlas represent specialized stem cell populations with gametogenic or somatic fates and provide a basis for deeper characterization of adult pluripotent stem cell populations in this colonial invertebrate.

## Results

### Identification of new markers from the two i-cell clusters of the ***Hydractinia* single-cell atlas**

The *Hydractinia* single-cell atlas was used to identify new markers from two i-cell clusters (Fig 2A) [43]. New markers were selected from the list of differentially expressed genes from these clusters based on two main criteria; those that had known stem cell expression or stem cell functions in other animals (*Pcna*, *Nop58*, *Mcm4*, *Ubr7*, *Uhrf1*) [16,45–48], or those that were highly expressed in one or the other i-cell clusters (*Pter, FoxQ2-like, Zcwpw1*) (S1 Table). The three different categories of markers, based on their expression in the *Hydractinia* single-cell atlas, are shown as Fig 2B: markers expressed in the i-cell cluster connected to somatic cell lineages (red), markers expressed in the i-cell cluster connected to the germ cell lineage (light blue), and markers expressed in both i-cell clusters (gold). The single-cell atlas expression patterns of the eight new markers are shown in Fig 2C, together with the expression pattern of *Piwi1*. Two markers are expressed in the somatic i-cell cluster (*Pter* and *FoxQ2-like*), one marker is expressed in the germ i-cell cluster (*Zcwpw1*), and five markers are expressed in both clusters (*Pcna*, *Nop58*, *Mcm4*, *Ubr7*, and *Uhrf1*). Gene orthology and nomenclature were based on reciprocal BLAST hits for all genes except *FoxQ2-like*, which could not be determined using this method. Instead, all *Hydractinia* Fox predicted protein sequences were combined with a previously published cnidarian forkhead DNA-binding domain dataset, and maximum likelihood phylogenetic analysis was performed [49]. The *Hydractinia Fox* gene that was characterized in this study grouped with the “*FoxQ2-like*” genes from both *Clytia hemisphaerica* and *Hydra vulgaris*, and thus has been named *FoxQ2-like* (S2 Fig).

### New markers have i-cell expression patterns in adult feeding polyps and are co-expressed with proliferating cells and *Piwi1*

To determine if the new marker genes were expressed in the i-cell band in adult feeding polyps, we performed RNA *in situ* hybridization using gene-specific riboprobes. Markers exclusively present in the i-cell cluster connected to somatic lineages are expressed in a band-like pattern in the epidermal layer of feeding polyps (*FoxQ2-like* and *Pter*, Fig 3Ai and 3Aii). The sizes of the cells vary from 5-8µm and are often in clusters of two or more (Fig 3Ai’ and Fig 3Aii’). In contrast, the marker unique to the i-cell cluster connected to the germ lineage showed no expression in feeding polyps (*Zcwpw1*, Fig 3Aiii and 3Aiii’). The markers that are expressed in both somatic and germ i-cell clusters have expression identical to the somatic i-cell cluster markers, with a characteristic band-like pattern in feeding polyps, in cells with 5-8µm diameter (*Pcna*, *Nop58*, *Mcm4*, *Ubr7*, and *Uhrf1*, Fig 3Aiv-3Aviii and Fig 3Aiv’-3Aviii’).

**Fig 3.**
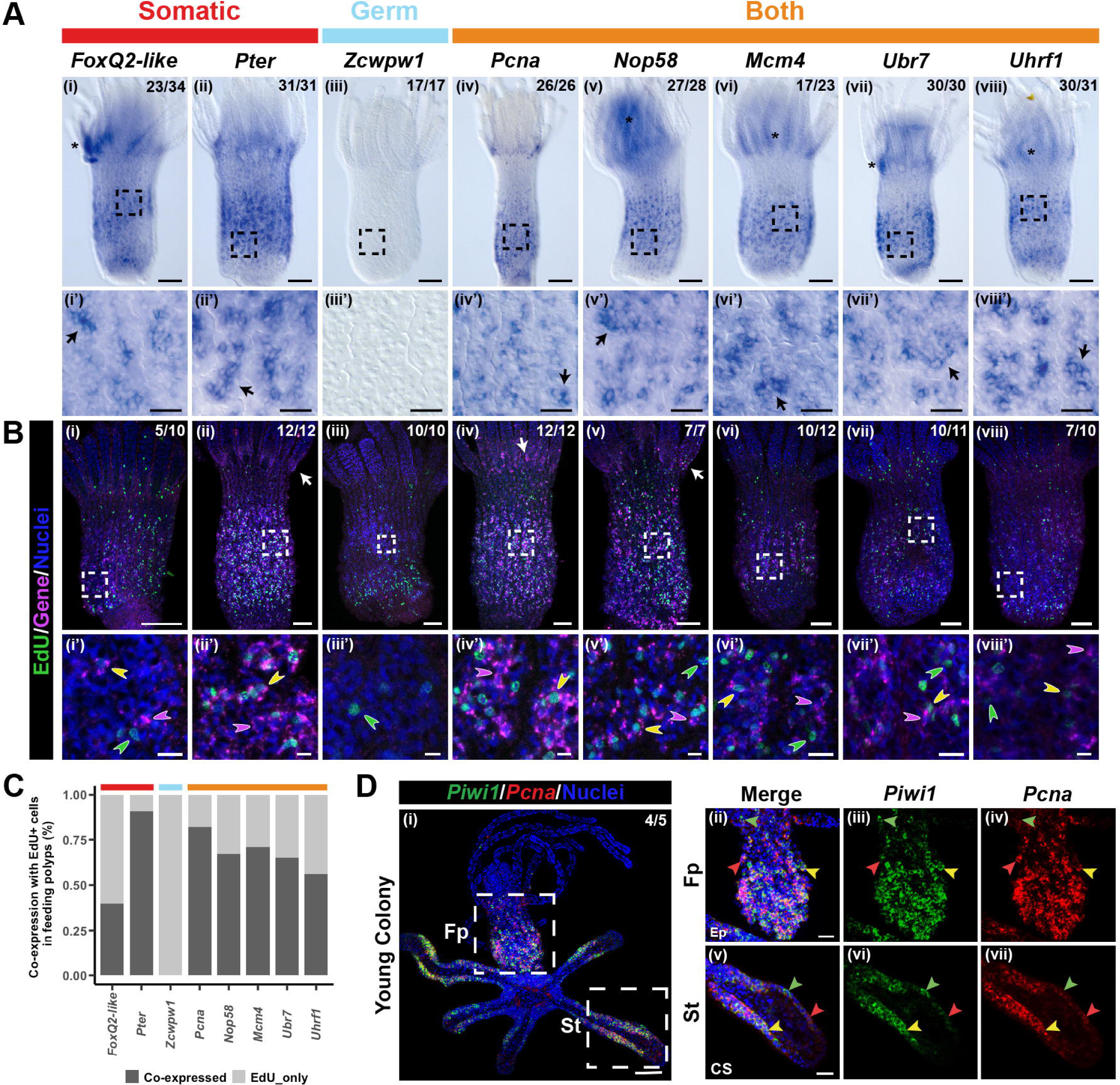
New marker genes are expressed in the feeding polyp i-cell band and are co-expressed in proliferating and *Piwi1^+^* cells. (A) Colorimetric *in situ* hybridization patterns of new markers in adult feeding polyps. Genes are organized according to the strategy used to identify them shown in Fig 2B and shown by colored bars above the panels; red indicates somatic i-cell genes, blue indicates the germ i-cell gene, and gold indicates the genes expressed in both somatic and germ i-cell clusters. Asterisks indicate background signal in the oral end of *Hydractinia* polyps that occasionally occurs during *in situ* hybridizations in *Hydractinia*. Black dashed boxes in the top panels indicate the region selected for higher magnification images in the bottom panels. Scale bars are 100µm in top panels (i) – (viii) and are 20µm in bottom panels (i’) – (viii’). Black arrows in the bottom panels indicate groups of cells. (B) Fluorescent *in situ* hybridization patterns of new markers co-labeled with EdU. Gene labels are the same as in (A). Green indicates EdU^+^ nuclei, pink indicates target gene expression, and blue is nuclei. White dashed boxes in the top panels indicate the region selected for higher magnification images in the bottom panels. Scale bars are 100µm in top panels (i) – (viii) and are 10µm in bottom panels (i’) – (viii’). White arrows indicate examples of target gene signal in the tentacles. Yellow arrowheads indicate target gene^+^/EdU^+^ cells, pink arrowheads indicate cells that only have expression of the gene of interest, and green arrowheads indicate cells that are positive for EdU only. (C) Bar chart showing the percent co-expression of a particular gene of interest and EdU in feeding polyps. Horizontal bars at the top are colored according to the strategy used to identify each new marker. (D) dFISH of *Piwi1* (green) and *Pcna* (red) in a young *Hydractinia* colony. (i) dFISH of a feeding polyp and stolon. Nuclei are blue. Numbers in the top right corner show the proportion of samples that reflect the image shown. Scale bar is 100µm. White dashed boxes in (i) indicate the higher magnification images shown in (ii) – (vii) which have 20µm scale bars. Fp is Feeding polyp body; St is stolon; Ep indicates ectodermal focal plane; CS indicates a cross-section focal plane. Yellow arrowheads indicate *Piwi1^+^*/*Pcna^+^* cells, green arrowheads indicate *Piwi1^+^* only expression, and red arrowheads indicate *Pcna^+^* only expression.

The EdU cell proliferation assay was used to determine if the new markers were expressed in proliferating cells in the feeding polyp. Markers expressed exclusively in the i-cell cluster linked to the somatic lineages displayed a wide range in the amount of overlap with EdU^+^ cells (*FoxQ2-like* and *Pter*, Fig 3Bi, 3Bi’, 3Bii, and 3Bii’). *FoxQ2-like* was expressed in fewer than half of EdU^+^ cells (10/25) while *Pter* was expressed in approximately 90% of EdU^+^ cells (42/46) (Fig 3C). The germ i-cell cluster marker*, Zcwpw1*, was not expressed in feeding polyps and therefore had no overlap with EdU^+^ cells (Fig 3Biii and 3Biii’). All five markers expressed in both i-cell clusters had a relatively high degree of overlap with EdU^+^ cells (*Pcna*, *Nop58*, *Mcm4*, *Ubr7*, and *Uhrf1*, Fig 3Biv-3Bviii and Fig 3Biv’-Bviii’’). Approximately 80% of EdU^+^ cells were also *Pcna*^+^ (69/84), while the degree of overlap for *Nop58* (55/82), *Mcm4* (47/66), *Ubr7* (35/54) and *Uhrf1* (20/36) was approximately 55-70% (Fig 3C). Examples of cells that were EdU^+^ only and marker gene^+^ only were present in all feeding polyps examined. There was also expression of several marker genes in cells at the base of the tentacles, most easily seen for *Pcna* (Fig 3Biv).

To investigate if the new markers were co-expressed with *Piwi1*, we first bioinformatically interrogated the *Hydractinia* single-cell atlas and assessed the percentage of *Piwi1^+^* cells that also expressed each of our new markers. We examined overlap separately in the somatic i-cell cluster, germ i-cell cluster, and both i-cell clusters (S2 Table). Marker genes in the somatic i-cell cluster (excluding *Zcwpw1*) had the lowest average co-expression with *Piwi1* at 22.6%, while the marker genes in the germ i-cell cluster (excluding *FoxQ2-like* and *Pter*) had the highest average co-expression with *Piwi1* at 47.8%, and marker genes present in both clusters had an average overlap of 29.2%. *Pcna* was chosen as a candidate gene to validate co-expression with *Piwi1* using double fluorescent *in situ* hybridization (dFISH), as it showed some of the highest overlap with *Piwi1* (35.3% across both clusters), and reliably gave good signal in *in situ* hybridization experiments. Young *Hydractinia* colonies grown on cover slips were used because this stage is more transparent and easier to image three-color fluorescence. Co-expression analysis of *Piwi1* and *Pcna* via dFISH showed overlap in cells in the stolons and the feeding polyp i-cell band (Fig 3D), though there are also clear examples of *Piwi1*^+^ only and *Pcna^+^* only cells (Fig 3Dii-3Dvii)). Of 353 randomly chosen cells expressing either gene, counted in four different young colonies, 32% co-expressed *Piwi1* and *Pcna*, 37% were *Piwi1^+^*only, and 31% were *Pcna^+^* only.

### New markers characterize different stages of sexual polyp sporosac development

The new marker genes show varied expression patterns in sexual polyps and their sporosacs. *FoxQ2-like*, a marker exclusive to the somatic i-cell cluster in the *Hydractinia* single-cell atlas, is not expressed in this polyp type (Fig 4Ai, 4Ai’ and 4Ai”). *Pter*, the other somatic i-cell cluster marker, is expressed only in the ectoderm of both the sexual polyp body and budding sporosacs (Fig 4Bi, 4Bi’ and 4Bi”). *Pter* appears to be expressed in a majority of cells in these regions, although individual cell morphology is unclear, making it difficult to identify cell types. In contrast, *Zcwpw1*, *Ubr7*, and *Uhrf1* are expressed in small to large sporosacs in the layer between the ectodermal and endodermal tissue where the gametes are produced (Fig 4Ci-4Ciii). The spatial location and morphology of the cells expressing these markers suggest that they are maturing spermatozoa [50], with sizes ranging between 3-4µm (Fig 4Ci’-4Ciii’ and Fig 4Ci’’-4Ciii’’). Only the outer layer of this compartment is stained, likely due to certain reagents in the *in situ* hybridization protocol not being able to penetrate deeper into the densely packed tissue. *Pcna*, *Nop58*, and *Mcm4* are expressed in small and budding sporosacs (Fig 4Di-4Diii). *Pcna* and *Mcm4* are expressed in cells with morphology similar to i-cells (Fig 4Di’ and 4Diii’, Fig 4Di’’ and 4Diii’’) while *Nop58*-expressing cells are similar to previously described somatic cells that encapsulate the maturing spermatogonia in the sporosacs (Fig 4Dii’ and 4Dii’’) [34]. The described *Piwi1* band-like pattern below the sporosacs in sexual polyps was not seen in any of the samples, likely due to the difficulty of including this region when dissecting this polyp type from the colony during sample preparation. Thus, we cannot assess whether the new marker genes are expressed in this region in sexual polyps in a similar manner to *Piwi1*.

**Fig 4.**
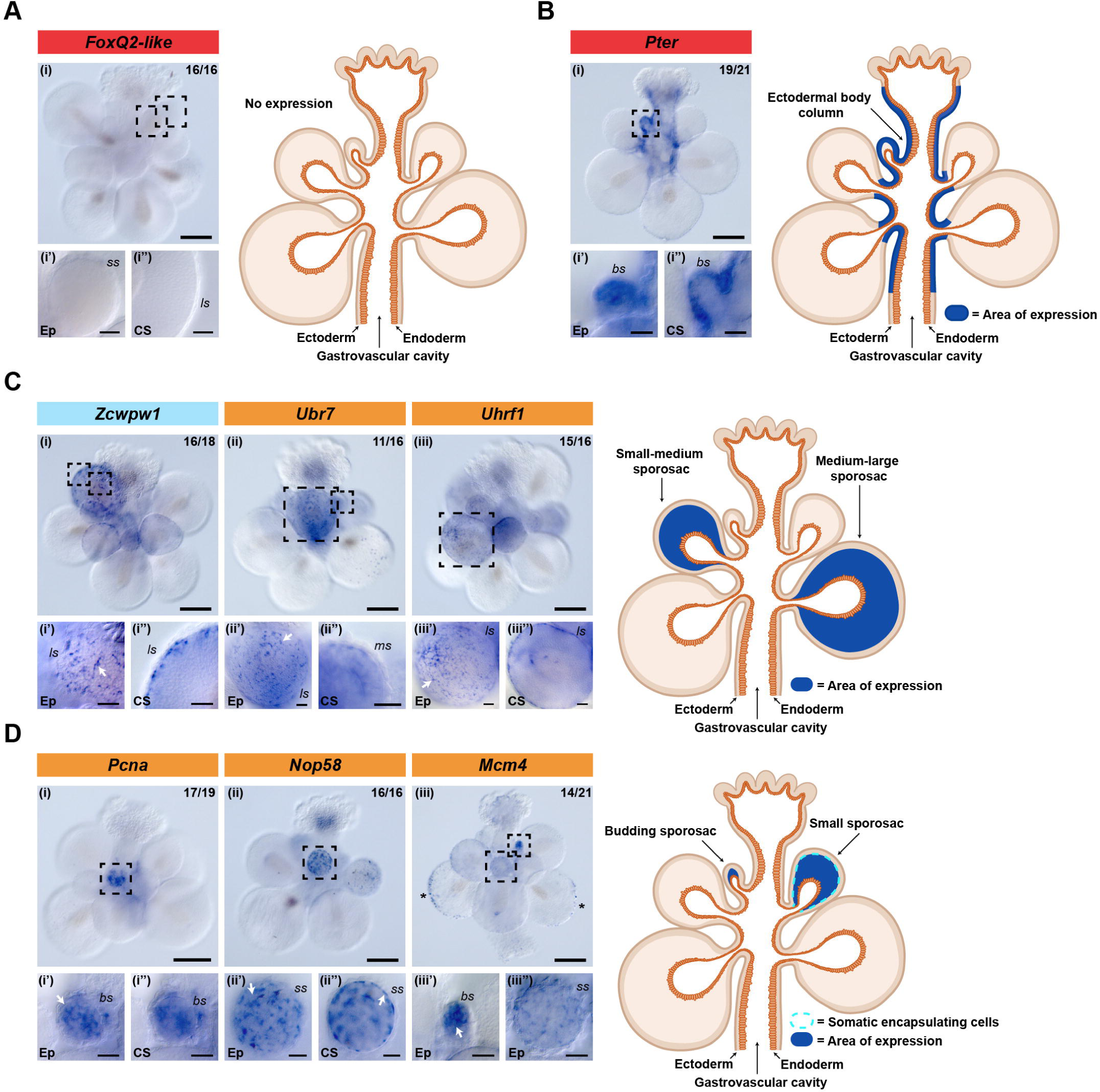
Expression patterns of new markers in male sexual polyps. Colorimetric *in situ* hybridization of new markers in male sexual polyps. Genes are organized according to the strategy used to identify them as shown in Fig 2B and indicated by colored bars above the panels; red indicates somatic i-cell genes, blue indicates the germ i-cell gene, and gold indicates the genes expressed in both somatic and germ i-cell clusters (A) *FoxQ2-like* (B) *Pter* (C) *Zcwpw1*, *Ubr7*, and *Uhrf1* (D) *Pcna*, *Nop58*, and *Mcm4*. The black dashed box in the upper *in situ* image indicates the region selected for higher magnification in the lower *in situ* images. Upper *in situ* images have 100µm scale bars and lower *in situ* images have 20µm scale bars. Numbers in the top right corner show the proportion of samples that reflect the image shown. White arrows indicate examples of target gene expression in different locations or cellular morphologies. Schematics on the right of each set of *in situ* hybridization panels summarize the expression pattern of each gene in male sexual polyps. Asterisks indicate a non-cellular background signal on large sporosacs (see *Mcm4*) that occasionally happens with *in situ* hybridization of *Hydractinia* tissue. Ep indicates an ectodermal focal plane; CS indicates a cross-section focal plane; bs labels budding sporosacs; ss labels small sporosacs; ms labels medium sporosacs; ls labels large sporosacs.

### New marker genes have dynamic expression during *Hydractinia* **feeding polyp head regeneration**

To determine if the new marker genes are involved in feeding polyp head regeneration, we decapitated feeding polyps, and examined their expression during wound healing, blastema formation, and tentacle bud regeneration stages (Fig 5A). Expression analysis during head regeneration shows that some genes are expressed in the blastema, while others are not. All maintained expression in the i-cell band during the time points analyzed (Fig 5B and Fig 5C, S3 Fig A and B). *Pcna*, *Mcm4*, *Pter*, and *Uhrf1* were expressed in the blastema tissue of the regenerating head at 24hpd, in cells with typical i-cell morphology (Fig 5Bii and 5Bii’, Fig 5Bv and 5Bv’, S3 Fig Aii and Aii’, S3 Fig Av and Avi). Expression of *Pcna*, *Mcm4*, *Pter*, and *Uhrf1* was also seen in the band-like region in wound healing, blastema, and tentacle bud regeneration timepoints, the same as in intact polyps (Fig 5Bi-5Bvi, S3 Fig Ai-Avi). The remaining marker genes (*FoxQ2-like*, *Nop58*, and *Ubr7),* were not expressed in the blastema (Fig 5Cii and 5Cv, S3 Fig Bv). They do, however, maintain signal in the i-cell band-like pattern throughout regeneration, similar to the other i-cell marker genes (Fig 5 Ci-Cvi, S3 Fig Biv-Bvi). *Zcwpw1* had no expression at any time point during head regeneration of feeding polyps, as expected (S3 Fig Bi-Biii and Bi’-Biii’). Aside from *Pcna*, no markers were expressed in the area around the wound during the wound-healing timepoint.

**Fig 5.**
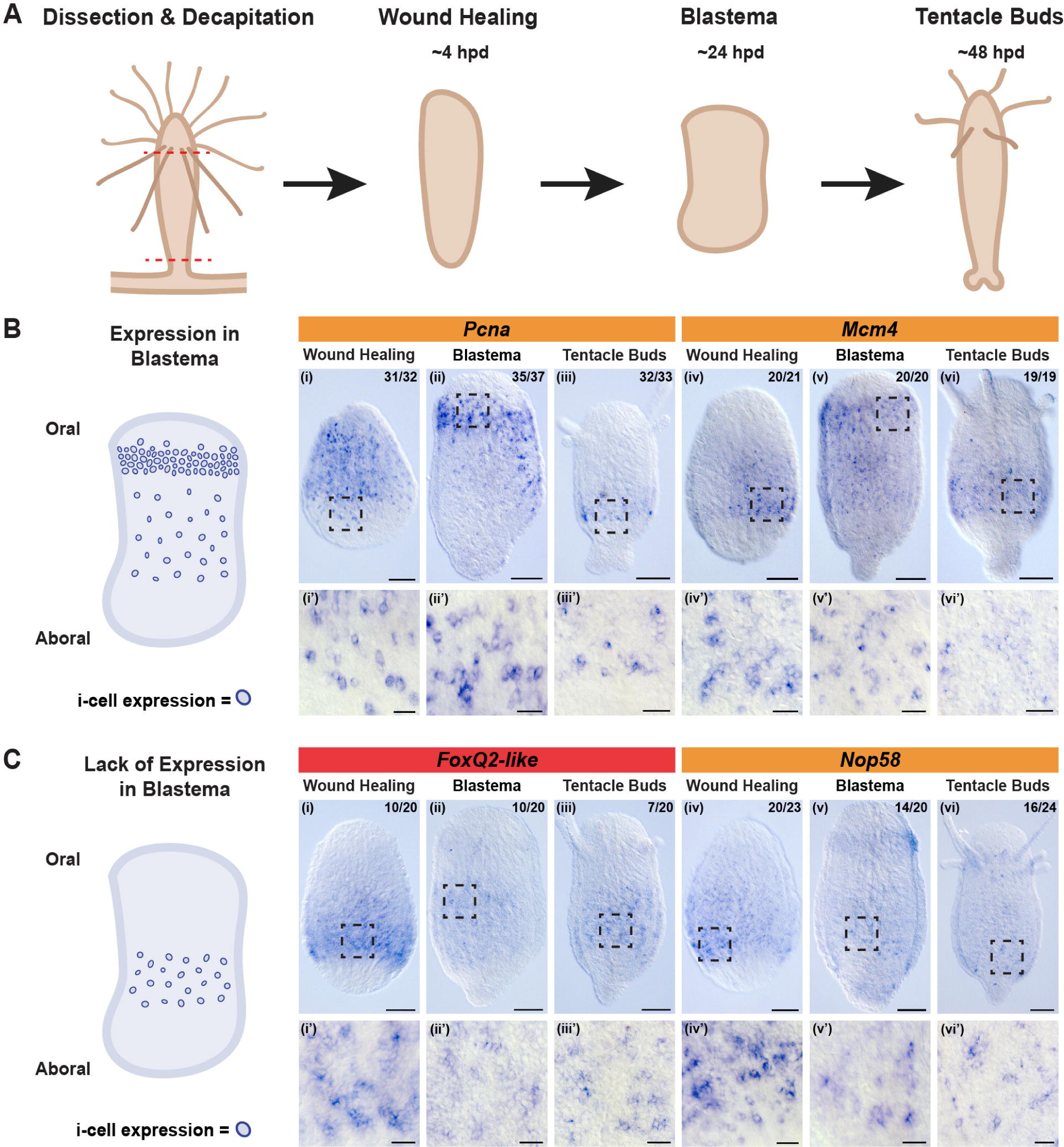
A subset of new markers are expressed in the blastema of the head during feeding polyp head regeneration. (A) Schematic showing the progression of *Hydractinia* feeding polyp head regeneration over a period of 48 hours post-decapitation (hpd). (B) Schematic of the blastema stage of regeneration depicting obvious expression in the blastema and the body column (left) and colorimetric *in situ* images (right) of *Pcna* and *Mcm4*. (C) Schematic of the blastema stage of regeneration depicting a lack of expression in the blastema (left) and *in situ* images (right) of *FoxQ2-like* and *Nop58*. The top panels in (B) and (C) have 100µm scale bars, bottom panels have 20µm scale bars. Black dashed boxes in (i) – (vi) indicate the higher magnification images shown in (i’) – (vi’). Numbers in the top right corner show the proportion of samples that reflect the image shown. Genes are organized by color according to the strategy used to identify them in Fig 2B; red indicates somatic i-cell genes, blue indicates the germ i-cell gene, and gold indicates genes expressed in both i-cell clusters.

### Consistent expression of new marker genes in stolon but not in all stages of developing feeding polyps in young colonies

We utilized young colonies (7-10 days post metamorphosis) to characterize the expression of the new marker genes and *Piwi1* in both stolon and multiple developing feeding polyp stages. We focused on the following stages of feeding polyps: (1) budding feeding polyps, (2) immature feeding polyps, and (3) mature feeding polyps. Except for *Zcwpw1,* all genes investigated were robustly expressed in the stolon (Fig 6 and S4 Fig). Expression of the genes in budding feeding polyps was also consistent, where *Piwi1*, *Pcna*, *Nop58*, *Pter*, and *Uhrf1* were ubiquitously expressed in nearly 100% of buds (Fig 6Ai-6Aiii and 6Ai’’-6Aiii’’, S4 Fig Aii, Av, Aii’’, Av’’). *Ubr7* had a slightly lower prevalence in budding polyps with ∼60% of samples showing expression (S4 Fig Aiv and Aiv’’). In contrast, *Mcm4* and *FoxQ2-like* expression was absent from most budding feeding polyps, with ∼90% of polyp buds lacking expression of either gene (Fig 6Aiv and 6Aiv’’, S4 Fig Ai and Ai’’).

**Fig 6.**
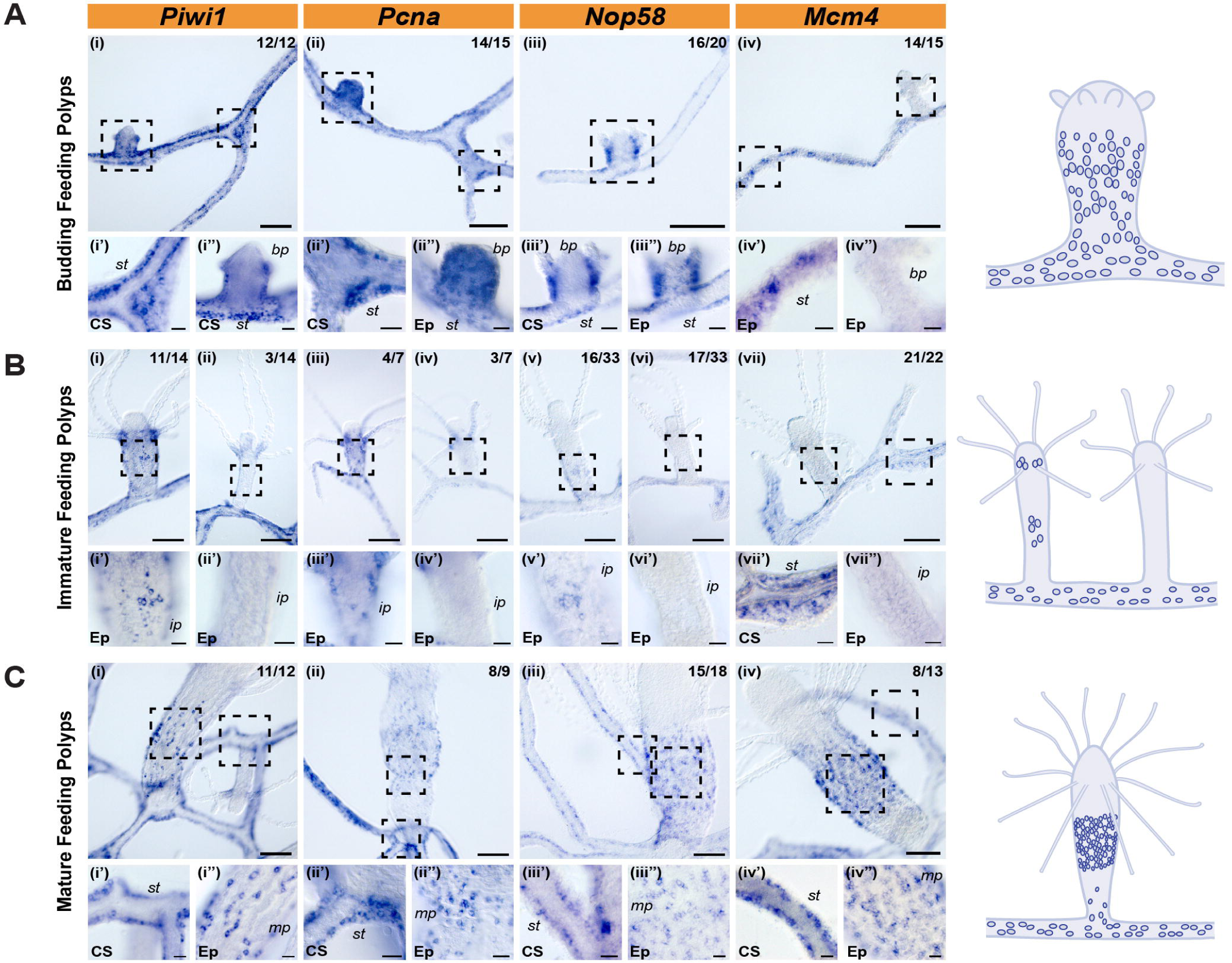
New marker genes are always expressed in the stolons of young colonies but are variably expressed in different developmental stages of feeding polyps. Colorimetric *in situ* hybridization of *Piwi1* and new marker genes *Pcna*, *Nop58*, and *Mcm4* in 7-10-day-old colonies focused on (A) budding polyps, (B) immature feeding polyps, and (C) mature feeding polyps. Genes are organized by color according to the strategy used to identify them in Fig 2B; gold indicates the genes expressed in both somatic and germ i-cell clusters. The top panels in (A), (B), and (C) have 100µm scale bars, while the bottom panels have 20µm scale bars. Black dashed boxes in (i) – (iv) indicate the higher magnification images shown in (i’) – (iv’) and (i’’) – (iv’’). Ep is the Ectodermal focal plane; CS is the cross-section focal plane; st labels stolon; bp labels budding polyps; ip labels immature polyps; mp labels mature polyps. Numbers in the top right corner show the proportion of samples that reflect the image shown for different developmental stages of feeding polyps. Schematics summarizing the observed expression of *Piwi1* and the new marker genes in different stages of feeding polyp development are shown to the right. Criteria for different stages of feeding polyps are as follows: budding feeding polyps are ∼100µm in length from the stolonal tissue and have between 0-4 budding tentacles; Immature feeding polyps are ∼300-400µm in length from the stolon and have between 6-8 tentacles that are about a body length in size; Mature feeding polyps are ∼600-800µm in length from the stolon and have 10 or more tentacles that are ∼1.5x their body length in size.

Expression of all genes in immature feeding polyps was variable. *Piwi1* was expressed in the ectodermal epithelia of 78.5% of immature polyps, including a high density of *Piwi1*^+^ cells at the base of the tentacles, but was excluded from the tentacles and hypostome (Fig 6Bi and 6Bi’). *Pcna*, *Nop58*, *FoxQ2-like*, *Pter*, and *Uhrf1* were expressed in 50-60% of immature feeding polyps, with the remaining polyps free of expression (Fig 6Biii-6Bvi and 6Biii’-6Bvi’, S4 Fig Bi-Biv, Bvii-Bviii, Bi’-Biv’, Bvii’-Bviii’). Each had a similar pattern to *Piwi1*, except for *Nop58* and *FoxQ2-like*, which seemed to be weakly expressed in the typical i-cell band-like pattern (Fig 6Bv and 6Bv’, S4 Fig Bi and Bi’). *Mcm4* and *Ubr7* expression was absent in most immature feeding polyps, with only 5-25% of polyps having signal (Fig 6Bvii and 6Bvii’’, S4 Fig Bvi and Bvi’’).

Finally, the signal for many genes in mature feeding polyps was nearly as robust as the expression in the stolonal tissue. *Piwi1*, *Pcna*, *Nop58*, *Pter*, and *Uhrf1* were expressed in 80-90% of mature feeding polyps in the typical i-cell band-like pattern seen in mature feeding polyps in adult colonies (Fig 6Ci-6Ciii and 6Ci’’-6Ciii’’, S4 Fig Ciii and Ciii’’, Cvi and Cvi’’). *Mcm4* and *Ubr7* had a lower percent expression, with 60-65%, but had the same i-cell band-like pattern (Fig 6Civ and 6Civ’’, S4 Fig Cv and Cv’’). *FoxQ2-like* had inconsistent expression in mature feeding polyps, with 30% of polyps having signal, and the remaining 70% devoid of signal (S4 Fig Ci-Cii and Ci’-Cii’). Of the mature feeding polyps that had *FoxQ2-like* signal, the pattern was present from the base of the polyp to the base of the tentacles, rather than the i-cell band-like pattern that was seen for *FoxQ2-like* in mature feeding polyps from adult colonies. Schematic representations of expression in stolonal tissue and different developmental stages of feeding polyps for all new markers and *Piwi1* in young colonies are shown on the right of Fig 6A-6C.

### Cycling cells are consistently present in the stolon and all feeding polyp developmental stages

To determine if the absence of expression of many of the marker genes in immature feeding polyps was simply due to a lack of proliferating i-cells and progenitors in this stage, we examined the expression of Histone *H1.1* in young colonies. *Hydractinia H1.1* is a member of the canonical histone family that is replication-dependent and expressed in the S-phase of cycling cells [50,51]. We found that *H1.1* is consistently expressed in all areas of young colonies; budding feeding polyps (Fig 7Ai and 7Ai’), immature feeding polyps (Fig 7Aii and 7Aii’’), mature feeding polyps (Fig 7Aiii and 7Aiii’’) and the stolon (Fig 7Ai’, 7Aii’, 7Aiii’), showing that proliferating cells are invariably present in each of these stages. In immature feeding polyps, *H1.1* signal was primarily located at a high density at the base of the tentacles, with some expression in the i-cell band (Fig 7Aii and 7Aii’’).

**Fig 7.**
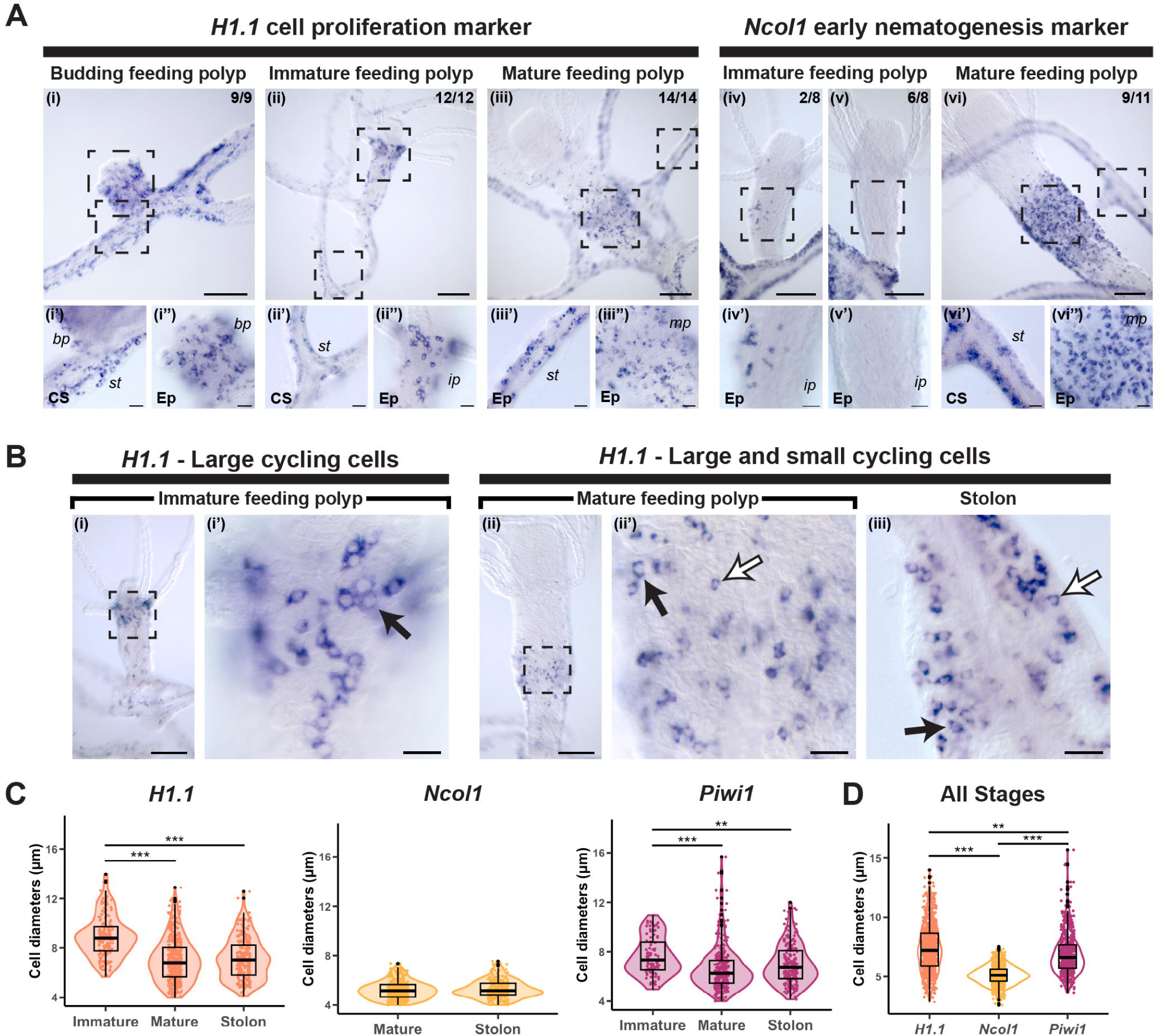
*H1.1^+^* cycling cells are present in all areas of the young colony and vary in size. (A) Colorimetric *in situ* hybridization of Histone *H1.1* (a marker of S-phase cycling cells) and *Ncol1* (a marker of non-cycling nematoblasts) in 7-10-day old young colonies of *Hydractinia*. Scale bars in top panels (i – vi) are 100µm, bottom panels (i’ – vi’’) are higher magnification images of the boxed region shown in top panels and have 20µm scale bars. Ep is the Ectodermal plane; CS is the cross-section plane; st labels stolon; bp labels budding polyps; ip labels immature polyps; mp labels mature polyps. Numbers in the top right corner show the proportion of samples that reflect the image shown for different developmental stages of feeding polyps. All samples showed gene expression in the stolonal tissue. (B) Comparison of *H1.1^+^*cycling cells found in immature feeding polyps (i, i’) and those in mature feeding polyps (ii, ii’) and stolonal tissue (iii). Rectangular panels (i, ii) have 100µm scale bars, and square panels (i’, ii’ and iii) have 20µm scale bars. Black dashed boxes in (i) – (ii) indicate the location of images in (i’) and (ii’). Black arrows show examples of large *H1.1^+^* cycling cells; white arrows with a black outline show examples of small *H1.1^+^* cycling cells. (C) Violin plots showing the cell diameters of *H1.1^+^*, *Ncol1^+^*, and *Piwi1^+^* cells across immature feeding polyps, mature feeding polyps, and stolons in young colonies. (D) Comparison of the diameters of all cells combined across immature feeding polyps, mature feeding polyps, and stolons for *H1.1*, *Ncol1*, and *Piwi1*. For (C) and (D), the significance bars at the top are from a post hoc TukeyHSD test and are labeled with the following significance codes: ** = p-adj<0.001, *** = p-adj<0.0001.

Next, we examined the expression of *Ncol1* in young colonies*. Ncol1* is a marker of committed nematocyte precursors, called nematoblasts, which are destined to become mature nematocytes in the colony [52]. The characteristic i-cell band-like pattern in mature feeding polyps is a well-known area of nematogenesis with strong *Ncol1* expression [29,30]. We found that *Ncol1* displays variation in expression patterns in developing polyps, most obvious when comparing immature feeding polyps and mature feeding polyps. *Ncol1* is expressed in only 25% of immature feeding polyps (Fig 7Aiv-v and 7Aiv’-v’) but is expressed in nearly 80% of mature feeding polyps (Fig 7Avi and 7Avi’’). *Ncol1* is consistently expressed throughout the stolon (Fig 7Aiv-v, 7Avi-vi’)

Additionally, *H1.1^+^, Ncol1^+^,* and *Piwi1^+^* cell diameters from different regions of young colonies were measured to further investigate the variation in expression seen in young colonies. Diameters of *H1.1*^+^ cells were measured across 11 stolonal sections (231 cells), 12 immature feeding polyps (150 cells), and 11 mature feeding polyps (385 cells). Similar polyp numbers, stolonal sections, and cell numbers were analyzed for *Ncol1* and *Piwi1*, except immature feeding polyps were excluded from *Ncol1* measurements as there were few polyps with signal, and those that did have signal had low numbers of *Ncol1*^+^ cells. One-way analysis of variance (ANOVA) was used to compare the mean cell size of gene^+^ cells between the different polyp stages and stolon for each gene and found that there was at least one statistically significant difference between means for *H1.1* (F(2, 763) = 75.21, p < .001) and *Piwi1* (F(2, 573) = 11.16, p < .001), but none for *Ncol1* (F(1, 644) = 1.73, p = 0.189). A post hoc Tukey test (TukeyHSD) was used to determine the significant comparisons from the *H1.1* and *Piwi1* ANOVA analyses. The size of *H1.1^+^* cells varied depending on the area of the colony in which they were located (Fig 7Bi-7Biii, Fig 7C). Significantly larger *H1.1*^+^ cells were found in immature feeding polyps (Fig 7Bi’), compared with mature feeding polyps or stolon, which both contained mixed populations of both large and small *H1.1*^+^ cells (Fig 7Bii’ and 7Biii, *H1.1*; p-adj<0.001 for both comparisons). Budding feeding polyps had a higher proportion of larger *H1.1*^+^ cells but this was less obvious than in the immature feeding polyp population (S5 Fig). In contrast, *Ncol1*^+^ cells have a relatively constant size in mature polyps and stolon (Fig 7Avi’ and 7Avii’’, Fig 7C). *Piwi1^+^* cells also showed differences in cell diameter depending on the region of the colony in which they reside; those in immature feeding polyps were significantly larger than those in either mature feeding polyps or stolon (Fig 7C) (*Piwi1*; immature-mature p-adj<0.001, immature-stolon p-adj=0.006). Cell diameters of *H1.1^+^*, *Ncol1^+^*, and *Piwi1^+^* cells in mature feeding polyps and stolonal tissue were not significantly different (Fig 7C) (*H1.1*; p-adj=0.2, *Ncol1*; p-adj=0.19, *Piwi1*; p-adj=0.11). Combined cell diameter measurements from immature feeding polyps, mature feeding polyps, and stolonal tissue of *H1.1^+^*, *Ncol1^+^*, and *Piwi1^+^* cells were also compared (Fig 7D). A one-way ANOVA found that there was at least one statistically significant difference in means (F(2, 1975) = 368.17, p < .001) and a post hoc TukeyHSD test showed that *H1.1^+^*cells were significantly larger than *Piwi1*^+^ cells (p-adj<0.001), and cells expressing either of these genes were significantly larger than *Ncol1^+^* cells (p-adj<0.001 for both comparisons).

Lastly, we performed dFISH using probes against *H1.1* and *Ncol1* to verify that there is no overlap between cycling cells (*H1.1*^+^ cells) and post-mitotic nematoblasts (*Ncol1*^+^ cells) and to further confirm that the absence of expression of *Ncol1* in many immature polyps is not due to an absence of proliferating cells (S6 Fig). All immature feeding polyps expressed *H1.1* while only one of these polyps also had *Ncol1* signal (S6 Fig i and i’). In contrast, both genes were expressed in all cases in mature feeding polyps and stolons with strong, non-overlapping signals (S6 Fig ii - iii and ii’ - iii’). Mature feeding polyps had the typical band-like pattern for both *H1.1* and *Ncol1* genes, with *H1.1* also expressed in cells beneath the tentacles, in the tentacles, and in the hypostome of the polyp (S6 Fig ii).

## Discussion

Here, we explore the two separate i-cell clusters from the *Hydractinia* single-cell atlas and show that they are biologically meaningful within the *Hydractinia* colony using a set of eight new marker genes. Our single germ i-cell marker (*Zcwpw1*) is exclusively expressed in gametogenic tissues in sexual polyps, our two somatic i-cell markers (*Pter* and *FoxQ2-like*) are expressed in somatic tissues, and our five markers present in both clusters (*Pcna*, *Nop58*, *Mcm4*, *Ubr7*, and *Uhrf1*) are expressed in somatic and gametogenic tissue throughout the colony. By exploring various biological contexts of the *Hydractinia* colony and examining the expression patterns of *Piwi1* and these eight new marker genes across different tissue types and during regeneration, we uncover heterogeneity in the spatial expression patterns of i-cell and progenitor markers that is less apparent when focused on a single tissue type or life history stage.

### Commonly studied i-cell and progenitor locations in the colony show minimal spatial expression variation of new markers

The i-cell band in the epidermis of the lower body column of the feeding polyp and stolonal tissue are known to harbor pluripotent adult stem cells and progenitors, and expression of genes in these areas is a well-established criterion for defining i-cell markers [2,7,26,29,39,40,53]. These areas also harbor both cycling and post-mitotic progenitor cells, however, and determining which cell type, cell subtype, or cell state a particular gene is expressed in is currently not feasible in this system, as markers have not been developed for each of these populations. All of the genes we characterized, except for the germ i-cell cluster marker, *Zcwpw1*, were consistently expressed in the i-cell band in adult feeding polyps and mature feeding polyps in a young colony, as well as throughout the epidermis of the stolon. Minimal variation amongst their patterns and cellular morphologies in these tissues makes it challenging to distinguish whether expression is in i-cells, progenitor cells, or both. In adult feeding polyps, all markers are expressed in a subset of cycling cells to varying degrees, similar to other i-cell markers in previous *Hydractinia* studies [29,39,54]. When combined with FISH, the presence of EdU^+^ only cells in the adult feeding polyps suggest the target genes (*FoxQ2-like*, *Pter*, *Pcna*, *Nop58*, *Mcm4*, *Ubr7*, and *Uhrf1I*) are not expressed in all proliferating cells. It is known that the feeding polyp band region has proliferating i-cells and nematocyte and neuronal progenitors, so it is possible our new markers are restricted to a subset of these cell types, though more evidence supporting this idea is needed. Bioinformatic analysis shows the number of cells co-expressing the new markers and *Piwi1* varies in the *Hydractinia* single-cell atlas i-cell clusters. Functional experiments like shRNA knockdown and subsequent quantification of differentiated cell types (e.g. nematocytes or neurons) will better resolve the potency of the cells expressing our new markers.

### Variable expression of new markers in different contexts provides a broader view of the i-cell population and hints at possible roles

The new markers had heterogeneous spatial expression patterns across different tissue types, different stages of polyp development, and during regeneration, and suggests a dynamic i-cell population. These results, based on gene expression in different locations in the colony and variable cellular morphologies, confirm the two i-cell clusters in the *Hydractinia* single-cell atlas are biologically meaningful. Additionally, the variations in patterns and cellular morphologies enabled us to hypothesize potential biological roles for these genes and compare them to known functions in other animals.

### Young colonies

Our results show that immature feeding polyps have particularly unique and variable patterns of expression of our new markers and *Piwi1*, where at least a subset of polyps show no expression. We reason that this is not due to technical issues, as evidenced by strong expression of the same gene in other areas of the same colony (e.g. stolon in Fig 6Bii *Piwi1*). This absence of expression was despite the presence of *H1.1*^+^ cycling cells in all immature feeding polyps examined. This is similar to what has been reported from another colonial hydrozoan species, the siphonophore *Nanomia bijuga*, where some areas with high cell proliferation were devoid of expression of *Piwi* and other i-cell markers [55]. It was hypothesized these cells were mitotically active epithelial cells or progenitor populations. In our case, the lack of expression of our i-cell and progenitor cell markers in cycling cells in immature polyps might be associated with a role in nematogenesis. We found that *Ncol1* expression – a marker of post-mitotic nematoblasts – was also absent in the majority of immature polyps, linking the presence of the typical i-cell band to nematogenesis. This raises the question of the identity of these cycling cells in immature polyps, given that they are unlikely to be nematoblasts and do not consistently express the known i-cell marker, *Piwi1*. The *H1.1^+^* cells and *Piwi1^+^* cells (when present) in immature polyps are on average significantly larger (average, 8.4µm; range in size, 4.0µm – 14.0µm; p-adj=0.006 or lower) compared to *H1.1^+^* and *Piwi1^+^* cells in mature polyps and stolon, where they vary in size from relatively small to large (average, 6.9µm; range in size, 2.9µm – 15.6µm). Different sizes of stem cells could indicate separate populations, and size has been shown to correlate with potency in other systems like humans and mice [56,57]. Based on previous descriptions of i-cells versus committed populations in *Hydra* [58], we hypothesize that the smaller sized proliferating cells (∼4-7µm) present in the mature polyp and stolon are committed somatic progenitors (e.g. cycling nematoblasts). All eight new markers, *Piwi1*, and *H1.1* are expressed in these smaller cycling cells, which are most frequently found in the i-cell band pattern or in the stolon. In contrast, the larger *H1.1^+^* and *Piwi1^+^*cells in immature polyps could represent the true pluripotent i-cells, as they are similar to the i-cell size range reported in the literature [27]. Alternately, these larger cycling cells might be committed progenitors of another somatic lineage, such as epithelial cells or gland cells. Further co-expression analyses with lineage-specific markers could help to reveal their identity.

### Sexual polyps

Expression of our markers in sexual polyps ranges from budding and small sporosacs (*Pcna, Nop58,* and *Mcm4*), to small-medium and medium-large sporosacs (*Zcwpw1, Ubr7,* and *Uhrf1*), to being either not expressed (*FoxQ2-like*) or expressed in regions outside the sporosacs (*Pter* and *Nop58*). None of the genes exactly match the expression pattern of *Piwi1*, which is expressed in various locations in sexual polyps, including the ectoderm and endoderm of the germinal zone, and within small-medium sporosacs (Fig 1C) [32]. It is likely that cells expressing our new markers that are within sporosacs in sexual polyps (i.e. *Zcwpw1*, *Ubr7*, *Uhrf1*, *Pcna*, *Mcm4*) do not represent pluripotent i-cells, but rather are populations committed to gametogenesis. Budding, small, and medium sporosacs contain germ cells that are in S-phase [50], and it is thought that all germ cells inside each sporosac in *Hydractinia* are synchronously developing [34]. Therefore, we hypothesize that genes expressed in budding or small sporosacs (*Pcna* and *Mcm4*) are functioning in the progression of S-phase before Meiosis I, whereas *Zcwpw1*, *Ubr7,* and *Uhrf1*, expressed in the small-medium, and medium-large sporosacs, are participating in Meiosis I or II. For *Zcwpw1*, this matches what is known in vertebrates, where it is a Meiosis I marker involved in the epigenetic regulation of cells during spermatogenesis [59,60]. Some other markers, namely *Pcna*, *Mcm4*, and *Uhrf1,* have also been shown to be present in cycling germ cells in other animals [61–65]. *Pter* is present only in the sexual polyp body ectoderm, suggesting it is involved in processes unrelated to gametogenesis. All marker genes are absent from the largest sporosacs, which previous studies have shown contain nearly mature or fully mature sperm ready to be released, based in part on the expression of *SLC9A10,* a marker of mature sperm cells [43].

### *Hydractinia* head regeneration

During *Hydractinia* head regeneration, we show that only four of our new markers (*Pcna*, *Mcm4*, *Pter*, and *Uhrf1*) have robust expression in the blastema of regenerating polyp heads, despite all genes (excluding *Zcwpw1*) exhibiting strong expression in the feeding polyp i-cell band. Expression in the blastema highlights markers that might play critical roles in head regeneration-specific i-cell proliferation or differentiation. The remaining markers (*FoxQ2-like*, *Nop58*, and *Ubr7*), while not expressed in the blastema at 24hpd, are expressed in the feeding polyp body column, revealing a previously unrecognized difference between the cells of the blastema and those in the i-cell band. What this difference means in a biological context is unclear. One possibility is that the blastema is responsible for making many more cell types, for example, the replacement of lost structural tissues, such as epithelial cells that form the hypostome or tentacles. In contrast, the cell types required for homeostasis of the mature feeding polyp might be expected to be predominantly nematocytes, which must be replaced after firing, therefore requiring nematoblasts and their progenitors to be a dominant cell type within the i-cell band. Recent findings from a study on tentacle regeneration in the medusa stage of the hydrozoan *Cladonema pacificum* support the idea that blastema i-cells and homeostatic i-cells represent heterogeneous stem cell populations [66]. In *Cladonema* medusa tentacle regeneration, the stem cells in the blastema primarily consist of repair-specific proliferative cells (RSPCs) that lack expression of classical i-cell markers like *Piwi* and do not appear to originate from the known resident homeostatic stem cells (RHSCs) in the tentacle bulb. The RSPCs primarily differentiate into the epithelial cells forming the regenerating tentacle while the RHSCs are believed to be multipotent stem cells that are focused on nematogenesis, similar to the resident tentacle bulb i-cells of the related hydrozoan, *Clytia* [67,68]. *Cladonema* medusa tentacle regeneration and *Hydractinia* polyp head regeneration exhibit some differences, with the i-cells in the regenerating head blastema of *Hydractinia* primarily originating from homeostatic stem cells in the i-cell band [29]. Both species, however, show relatively little expression of *Piwi1* in the blastema region, and knockdown of *Piwi1* in *Hydractinia* does not inhibit cell proliferation in the blastema [29]. This suggests that like *Cladonema*, the blastema i-cells of *Hydractinia* may be a proliferative subtype primarily responsible for replacing missing epithelial tissue, whereas the i-cells in the feeding polyp band likely represent a homeostatic population focused on nematogenesis. Taken together, both systems appear to have transcriptionally distinct i-cells in the regenerating blastema compared to their homeostatic populations found in the i-cell band or tentacle bulb, with the variation in expression of our new marker genes reflecting this heterogeneity.

### I-cells and progenitors display dynamic gene expression patterns depending on the context

We demonstrate that the i-cells and progenitors in *Hydractinia* are transcriptionally dynamic by characterizing the expression of eight new marker genes in different biological contexts throughout the colony and during polyp head regeneration (Fig 8). We show the two i-cell clusters in the *Hydractinia* single-cell atlas represent real biological heterogeneity, likely being a mixture of pluripotent stem cells and multipotent progenitor cells responsible for diverse processes with at least two distinguishable subsets: those involved in contributing to somatic lineages, and those involved in providing the germ cell lineage. These results are similar to recent work in acoels, planarians, annelids, and the jellyfish *Cladonema*, where adult stem cells exhibit transcriptional heterogeneity rather than being a molecularly homogeneous population [3,5,15,66]. In conclusion, we provide eight new markers of i-cells and progenitors in *Hydractinia*, demonstrate substantial heterogeneity in the spatial expression patterns of the new markers and *Piwi1*, and highlight the importance of examining multiple biological contexts to better understand cell clusters and genetic markers obtained from single-cell expression analyses, and i-cells in general.

**Fig 8.**
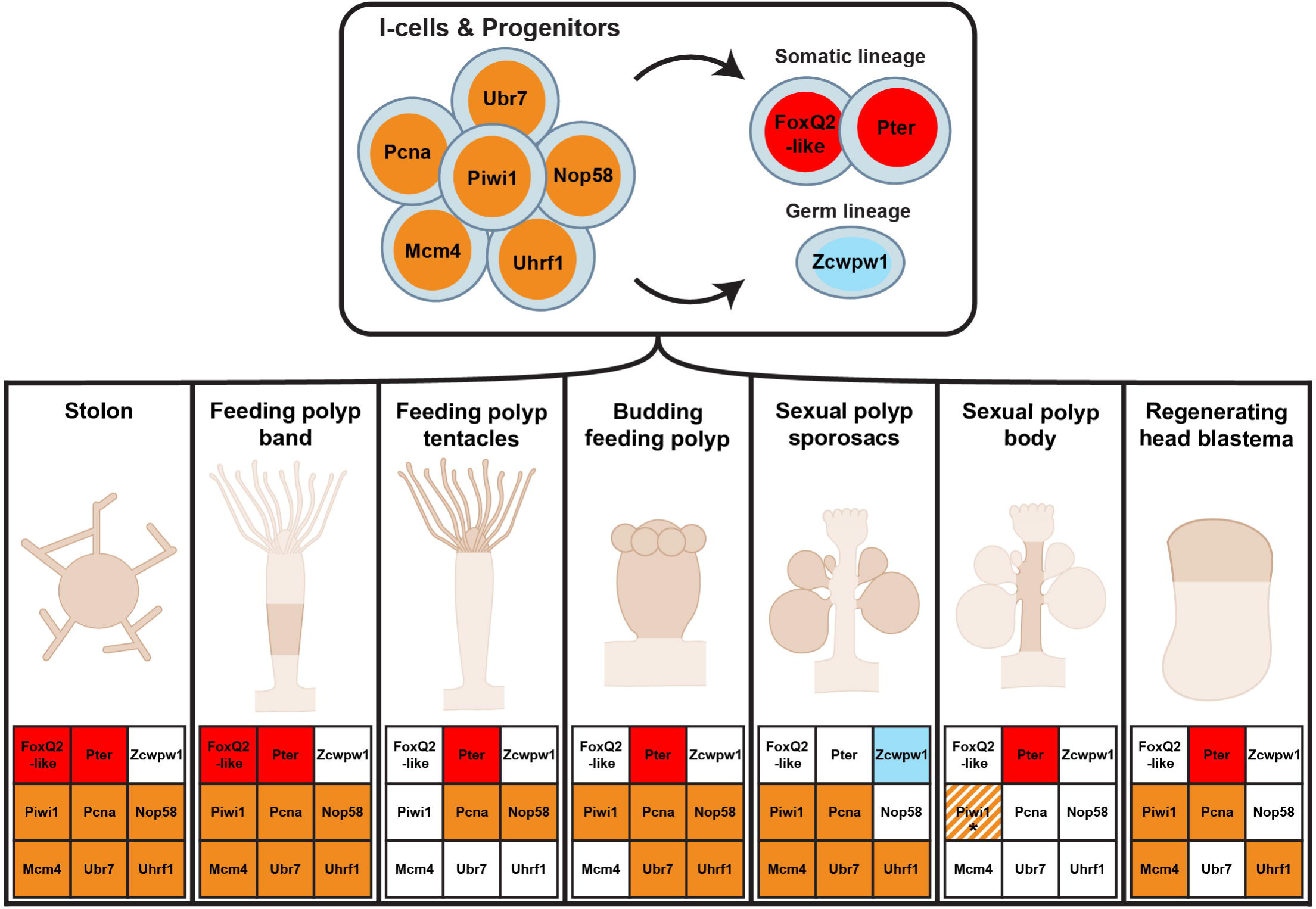
A model for understanding i-cells in a colonial animal. Putative cell lineages of new markers; somatic lineage (*FoxQ2-like* and *Pter*); germ cell lineage (*Zcwpw1*); pluripotent i-cells and their progenitors (*Piwi1*, *Mcm4*, *Pcna*, *Uhrf1*, *Nop58*, and *Ubr7*). Schematics and grids at the bottom summarize the expression of each new marker in different contexts, where a filled, colored box indicates gene expression is present, and an unfilled box indicates expression is absent. The striped colored box with an asterisk beneath *Piwi1* in the sexual polyp body section indicates that this gene has expression in parts of the sexual polyp body but differs from the patterns obtained using the new somatic lineage marker *Pter*.

## Materials and methods

### Animal husbandry and collection

*Hydractinia symbiolongicarpus* colonies were cultured and fed as previously described [39,69]. Young colonies were grown by inducing three-day-old larvae to metamorphose by incubating them in 116mM CsCl solution in 30 ppt filtered seawater (FSW) for 3 hours. After incubation, metamorphosing larvae were rinsed twice in FSW and settled onto 15mm round coverslips (CellTreat # 229172) and placed into 4- or 24-well plates (CellTreat # 229103 and Corning # 07-200-84). Animals were fed small or mashed artemia (SEP-ART GSL) daily, starting two days post-metamorphosis. The solution in each well was changed every 1-2 days with FSW containing 60□μg/mL penicillin and 50□μg/mL streptomycin (Sigma-Aldrich # P3032 and #S6501).

### Feeding polyp head regeneration

Adult colonies were relaxed in 4% MgCl_2_ consisting of 50% distilled water and 50% FSW for 10-15 minutes. Once relaxed, adult feeding polyps were cut from the stolonal mat using Vannas-type micro scissors (TedPella #1341). Dissected polyps were then decapitated just aboral to the tentacles using a Feather™ microscalpel (Electron Microscopy Services #72045-45). Animals were left in 90mm glass petri dishes with FSW to regenerate before fixation at appropriate time points (see below for *Hydractinia* fixation). FSW was changed daily until fixation.

### Fixation and *in situ* Hybridization

Animals were relaxed in 4% MgCl_2_ consisting of 50% distilled water and 50% FSW for 10-15 minutes and then fixed for 90 seconds in Fix 1 (0.2% glutaraldehyde, 4% paraformaldehyde, 0.1%Tween20 in FSW). Fix 1 was removed and samples were incubated in ice-cold Fix 2 (4% paraformaldehyde, 0.1% Tween20 in FSW), with rocking at 4°C for 90 minutes. Following fixation, samples were washed with ice-cold PTw (1x phosphate-buffered saline (PBS) with 0.1% Tween20). They were then dehydrated in increasing concentrations of methanol in PTw (25%, 50%, 75% and 100%). Samples were stored at -20°C for at least 24 hours.

Digoxigenin (DIG)-labeled and/or Fluoresecein-labeled riboprobes for selected genes were generated with the SP6 or T7 MEGAscript kit (Cat #AM1334, #AM1330, Ambion, Inc., Austin, TX, USA) using the open reading frames (ORFs) of cloned *Hydractinia* genes as template and following manufacturer’s recommendations. Accession numbers for each gene can be found in S3 Table. Primer sequences used to amplify genes from cDNA for cloning can be found in S4 Table.

### Fluorescent *in situ* hybridization

Samples were rehydrated with decreasing concentrations of methanol in PTw, followed by several washes in PTw. Samples were then washed for five minutes each in 1% triethylamine in PTw (TEA), 0.6% acetic anhydride in TEA, and 1.2% acetic anhydride in TEA, followed by several washes in PTw. Samples were pre-hybridized for 2-4 hours at 55°C in hybridization buffer (4M urea, 0.1 mg/ml yeast tRNA, 0.05 mg/ml Heparin, 5x SCC pH7.0, 0.1% Tween20, 1% SDS in DEPC-treated H2O). DIG-labeled probes were diluted to a concentration of 1-2 ng/ul in hybridization buffer, preheated to 90°C for 10 minutes, before being added to samples and left to hybridize for ∼40 hours at 55°C.

After hybridization, animals were washed once in hybridization buffer at 55°C for 40 minutes and then subjected to a series of post-hybridization washes with decreasing hybridization buffer concentrations in 2x SSC (at 55°C), followed by washes with decreasing concentrations of 0.2x SSC in PTw, and finally washes in PTw at room temperature (RT). Endogenous peroxidase activity was quenched by two 30-minute washes in 3% hydrogen peroxide (H_2_O_2_), followed by further washes in PTw. Two 10-minute washes in maleic acid buffer (MAB, 100mM Maleic acid, 150mM NaCl, pH7.5) were then conducted. Samples were then blocked for one hour in Blocking Buffer (Sigma-Aldrich 11096176001 diluted 1:10 in MAB). The probe was detected by incubating samples overnight at 4°C with a 1:1500 dilution of Anti-DIG-POD antibody (Roche, Catalog# 11207733910) in Blocking Buffer. Unbound antibody was removed by washing samples several times at room temperature in MABX (MAB containing 0.1% Triton X-100). To develop fluorescent signal, samples were incubated in Tyramide development solution (2% Dextran sulfate, 0.0015% hydrogen peroxide, 0.2mg/ml Iodophenol, 1:100 Alexa Fluor 594 Tyramide Reagent (Thermo Scientific, Cat. # B40957) in PTw) for eight minutes, and then washed several times in PTw. Nuclei were stained using Hoechst dye 33342 (ThermoFisher # H1399), and samples were left in Fluoromount (Sigma-Aldrich # F4680) for 1-3 days before mounting and imaging with a Zeiss LSM 710 confocal microscope.

### Double fluorescent *in situ* hybridization

Samples were fixed, stored, and treated as detailed above for fluorescent *in situ* hybridization, however, dFISH required both a fluorescein-labeled probe and a DIG-labeled probe to be added and incubated together at the hybridization step. Following washes and the anti-DIG-POD development reaction using Alexa Fluor 594 Tyramide Reagent, samples were incubated in 100 mM glycine stock solution pH 2.0 for 10 min at RT, and then washed five times in PTw for 5 minutes each. They were then washed in MABX two times for 10 minutes each and incubated in Blocking Buffer as above. The fluorescein-labeled probe was detected using an Anti-Fluor-POD antibody (Roche, Catalog# 11426346910), diluted 1:1500 in Blocking Buffer. For the second fluorescent reaction, samples were incubated in Tyramide development solution (2% Dextran sulfate, 0.0015% hydrogen peroxide, 0.2mg/ml Iodophenol, 1:100 Alexa Fluor 488 Tyramide (ThermoFisher # B40953) in PTw) for eight minutes, and then washed several times in PTw. Nuclei were stained using Hoechst dye 33342 (ThermoFisher # H1399), and samples were left in Fluoromount (Sigma-Aldrich # F4680) for 1-3 days before mounting and imaging with a Zeiss LSM 710 confocal microscope.

### Colorimetric *in situ* hybridization

After fixation, samples were rehydrated with decreasing concentrations of methanol in PTw, followed by several washes in PTw, before being heated to 85°C for 20 minutes to inactivate endogenous alkaline phosphatases. Samples were then washed for five minutes each in 1% triethylamine in PTw (TEA), 0.6% acetic anhydride in TEA, and 1.2% acetic anhydride in TEA, followed by several washes in PTw. Samples were then pre-hybridized for 2-4 hours at 55°C in hybridization buffer (4M urea, 0.1 mg/ml yeast tRNA, 0.05 mg/ml Heparin, 5x SCC pH7.0, 0.1% Tween20, 1% SDS in DEPC-treated H_2_O). DIG-labeled probes were diluted to a concentration of 1-2 ng/ul in hybridization buffer, preheated to 90°C for 10 minutes, before being added to samples and left to hybridize for ∼40 hours at 55°C. After hybridization, animals were washed once in hybridization buffer at 55°C for 40 minutes, and then through a series of post-hybridization washes with decreasing hybridization buffer concentration in 2x SSC (at 55°C), followed by washes with decreasing concentrations of 0.2x SSC in PTw at room temperature, and finally washes in PTw at room temperature. Two 10-minute washes in maleic acid buffer containing 0.1% Triton X-100 (MABX) were then conducted before samples were blocked in Blocking Buffer (see above), in MAB for at least one hour at room temperature. Probes were detected by incubation overnight at 4°C in a 1:5000 dilution of Anti-DIG-AP antibody (Roche, Catalog# 11093274910) in Blocking Buffer. Samples were then washed in MABX six times and then washed several times in alkaline phosphatase (AP) buffer (100 mM NaCl, 50 mM MgCl_2_, 100 mM Tris -pH 9.5, and 0.5% Tween20 in H_2_O). Finally, they were incubated in AP buffer containing 0.33 mg ml^−1^ NBT and 0.165 mg ml^−1^ BCIP. The solution was refreshed every 2 hours until the development reaction had proceeded to the desired point. The development reaction was stopped by washing several times in PTw and specimens were cleared using the EtOH clearing protocol previously described for *Hydra* [21]. Finally, samples were mounted in 80% glycerol in PBS and imaged using a Zeiss Imager.M2 compound light microscope.

### EdU assay and quantification

EdU experiments were performed as previously described [39]. Samples were incubated in EdU for 30 minutes before fixation and methanol dehydration. The Click-iT EdU (Invitrogen, Catalog# C10340) detection reaction was carried out for 1 hour at room temperature following the manufacturer’s recommendations. When combined with fluorescent *in situ* hybridization, the Click-iT EdU detection reaction was performed at the end of the protocol before nuclei staining.

Fiji was used to analyze confocal images of feeding polyps used for the FISH + EdU assay [70]. EdU, gene, and nuclei channels were separated and enhanced using the brightness and contrast option. Firstly, each EdU^+^ cell was outlined using the circle and draw options. This channel was then overlayed with the gene and nuclei channels to quantify the number of cells that had overlapping EdU and gene signals. All cells that were EdU^+^ only were annotated with squares using the draw option and then counted.

### Single-cell analysis

Single-cell data and analysis was obtained from Schnitzler et al., 2024 and the *Hydractinia* Genome Project Portal (https://research.nhgri.nih.gov/hydractinia/). Co-expression percentages of *Piwi1* and our new markers from S2 Table were obtained using the single-cell browser from the *Hydractinia* Genome Project Portal (see “gene coexpression” option). Single-cell plots were made using the R ‘Seurat’ package [71].

### BLAST orthology analysis and Fox gene ML phylogeny

A reciprocal BLASTP was done on the new markers in this study to verify their orthology. A BLAST E-value cutoff of 10^-100^ was used to determine whether genes needed further verification via gene trees. The *Fox-Q2-like* gene was the only gene that could not be resolved via BLAST. To generate the Fox tree shown in S2 Fig, a total of 155 full-length protein-coding sequences were aligned. The Fox protein alignment file from [49] which included only the forkhead domain of each protein was used, and we added 12 *Hydractinia* sequences to this alignment. We renamed the sequences for clarity and consistency (S5 Table). The final alignment file is provided in the supplementary material (S1 File). ProtTest 3 was used to select the best-fit model of protein evolution for the alignment, which was PROTGAMMALG (‘LG’ indicates the substitution matrix, and ‘gamma’ specifies gamma-distributed rates across sites). Maximum-likelihood (ML) analyses were performed using RaxML v. 8.2.11 [72]. ML branch supports are rapid RAxML Bootstrap values (500 replicates). The resulting tree was displayed with FigTree v.1.4.4 (http://tree.bio.ed.ac.uk/software/figtree/) and annotated in Adobe Illustrator®.

### Image processing and analysis

Zeiss LSM 710 confocal images were processed using Fiji and Z-stack maximum likelihood projections [73]. Projections made from 20x images used all slices within a stack, while 40x images typically used 2-5 slices. The signal was enhanced using the brightness and contrast options. Zeiss Imager.M2 images were processed using Adobe Photoshop® software. The following options were used to enhance images: levels, hue, color balance, and photo filters.

Fiji was used to measure cell diameters from various colorimetric *in situ* hybridization images [70]. Results were put into an Excel sheet and uploaded to R version 4.2.1 for statistical analysis and data visualization using the R ‘stats’ and ‘ggplot’ packages [74]. All figures were made using Adobe Illustrator® software.

## Acknowledgments

We want to thank Amanda Yeo, Isabella Cisneros, and Dr. Gonzalo Quiroga-Artigas for their assistance in preliminary data collection. Thank you to members of the Schnitzler lab and Martindale lab for their discussion and insights. We would also like to thank the Martindale lab for allowing us to use their microscopy equipment.

## Author contributions

J.W, C.S, and D.J conceived and designed the study. J.W, C.S, and D.J wrote the paper. J.W generated single-cell plots and analysis of co-expression of genes in this dataset. J.W and D.J performed ISH, FISH, EdU, and dFISH experiments. J.W and D.J performed colorimetric and confocal imaging. J.W performed image analysis, EdU quantification, and cell measurements. J.W created figures and schematics. C.S acquired funding.

## Supporting information

**S1 Fig. Expression patterns of several stem cell genes in the *Hydractinia* single-cell atlas.** These genes have been reported previously as having i-cell expression patterns in *Hydractinia* [29,31,35,39,75]. None of these genes had patterns in the atlas that were restricted to just the two i-cell clusters.

**S2 Fig. Maximum likelihood phylogenetic tree of Fox-like genes in *Hydractinia*.** The best fit model was PROTGAMMALG. Maximum-likelihood analysis was done using RAxML. The tree is unrooted and consists of non-metazoan and metazoan forkhead domains. The nodes are labeled with rapid RAxML Bootstrap values (500 replicates). Only Bootstrap percentages >50% for nodes outside Fox families and the group of sequences related to the *Hydractinia FoxQ2-like* gene are labeled. The asterisk indicates the *FoxQ2-like* gene investigated in this paper.

**S3 Fig. Additional new marker expression patterns at three time points of feeding polyp head regeneration.** (A) Schematic of the blastema stage of regeneration depicting obvious expression in the blastema and the body column (left) and colorimetric *in situ* images (right) of new markers *Pter* and *Uhrf1*. (B) Schematic of the blastema stage of regeneration depicting a lack of expression in the blastema (left) and colorimetric *in situ* images (right) of new markers *Zcwpw1* and *Ubr7*. The top panels in (A) and (B) have 100µm scale bars, bottom panels in (A) and (B) have 20µm scale bars. Black dashed boxes in (i) – (vi) indicate the higher magnification images shown in (i’) – (vi’). Numbers in the top right corner show the proportion of samples that reflect the image shown. Genes are organized by color according to the strategy used to identify them in Fig 2B; red indicates somatic i-cell genes, blue indicates the germ i-cell gene, and gold indicates the genes expressed in both somatic and germ i-cell clusters.

**S4 Fig. Additional new marker genes are always expressed in the stolons of young colonies but are variably expressed in different developmental stages of feeding polyps.** Colorimetric *in situ* hybridization images of 7-10-day-old young colonies for new marker genes *FoxQ2-like*, *Pter*, *Zcwpw1*, *Ubr7* and *Uhrf1*. Genes are organized by color according to the strategy used to identify them in Fig 2B; red indicates somatic i-cell genes, blue indicates the germ i-cell gene, and gold indicates the genes expressed in both somatic and germ i-cell clusters. The top panels in (A), (B), and (C) have 100µm scale bars, while the bottom panels in (A), (B) and (C) have 20µm scale bars. Black dashed boxes in (i) – (viii) indicate the higher magnification images shown in (i’) – (viii’) and (i’’) – (vi’’). Ep is the Ectodermal focal plane; CS is the cross-section focal plane; st labels stolon; bp labels budding polyps; ip labels immature polyps; mp labels mature polyps. Numbers in the top right corner show the proportion of samples that reflect the image shown.

**S5 Fig. Diameters of *H1.1^+^* cells across all feeding polyp developmental stages and stolon in young colonies.** Violin plots showing the cell diameters of *H1.1^+^* cells across budding feeding polyps, immature feeding polyps, mature feeding polyps, and stolons in young colonies. Significance bars at the top are labeled with the following significance code: *** = p-adj<0.0001.

**S6 Fig. *H1.1* cycling cells are present in immature feeding polyps despite an absence of *Ncol1* signal.** dFISH of *H1.1* and *Ncol1* in young colonies. Immature feeding polyp (i, i’), mature feeding polyp (ii, ii’), and stolon (iii, iii’). Red indicates *H1.1* expression, Green indicates *Ncol1* expression, and Blue indicates nuclei. Numbers in the top right corner (i) – (iii) show the proportion of samples that reflect the image shown and are colored corresponding to *H1.1* (red) and *Ncol1* (green). Scale bars in top panels (i) – (iii) are 100µm, bottom panels (i’) – (iii’) are of the white boxed region shown in top panels and have 20µm scale bars. White arrows indicate *H1.1^+^* cells at the base of the tentacles and in the hypostome. Ep is the Ectodermal focal plane; CS is the cross-section focal plane; st labels stolon; ip labels an immature polyp; mp labels a mature polyp.

**S1 Table. Pannzer descriptions of new markers from i-cell clusters.** Pannzer-annotated molecular and biological functions of new markers from i-cell clusters from the *Hydractinia* single-cell atlas [43].

**S2 Table. Percent overlap of *Piwi1* with new markers in i-cell clusters from the *Hydractinia* single-cell atlas.** Percent overlap of *Piwi1* with new markers from i-cell clusters in the *Hydractinia* single-cell atlas [43]. Overlap is examined in the somatic i-cell cluster, germ i-cell cluster, and both i-cell clusters. For example, the percent co-expression of *Piwi1* and *Pter* was 33.1% in the somatic i-cell cluster (119/360), 20.6% in the germ i-cell cluster (74/359), and 26.8% in both i-cell clusters (193/719).

**S3 Table. Accession numbers for all genes investigated in this study.**

**S4 Table. Primer sequences for cloning of all genes investigated in this study.**

**S5 Table. Original and renamed Fox domain-containing sequences.** A list of all genes from [49] that were renamed for clarity in the Fox gene ML phylogeny.

**S1 File. Fox gene alignment.** Fasta-formatted file containing the Fox protein alignment used in constructing the Fox gene ML phylogeny.

