## Supplementary figures and images for "Characterization of eight new *Hydractinia* i-cell markers reveals underlying heterogeneity in the adult pluripotent stem cell population"

### S1 Fig

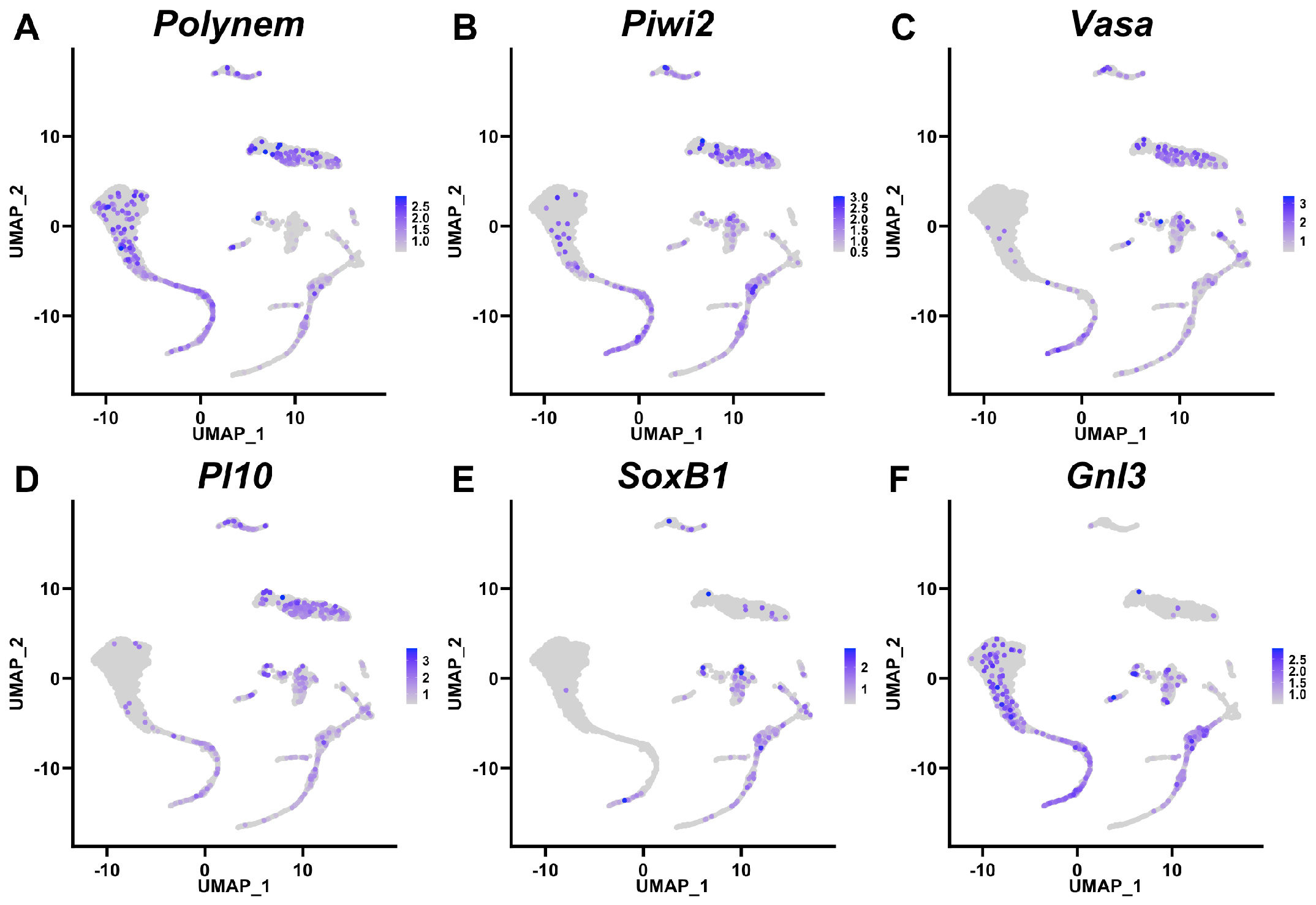

### S2 Fig

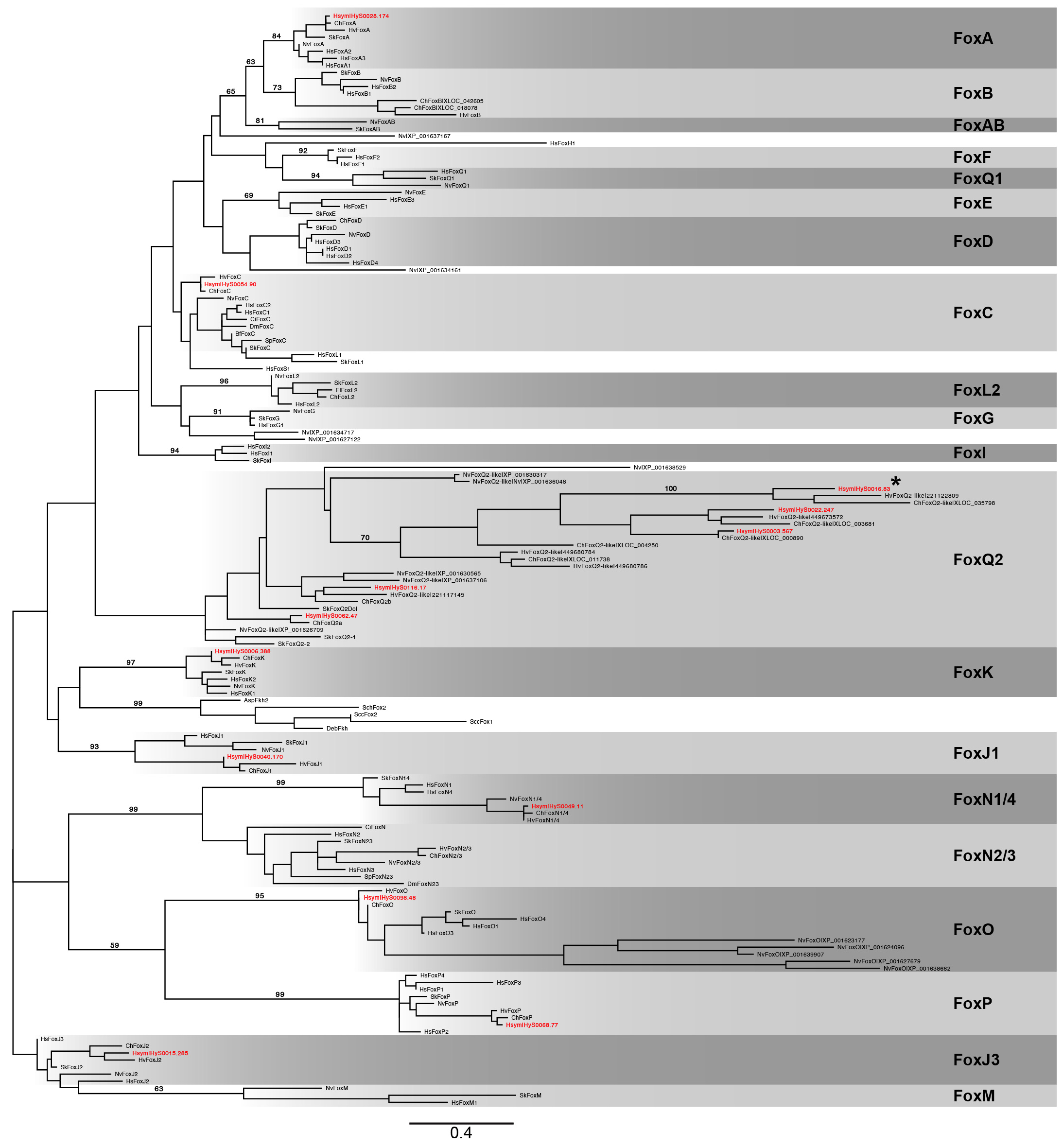

### S3 Fig

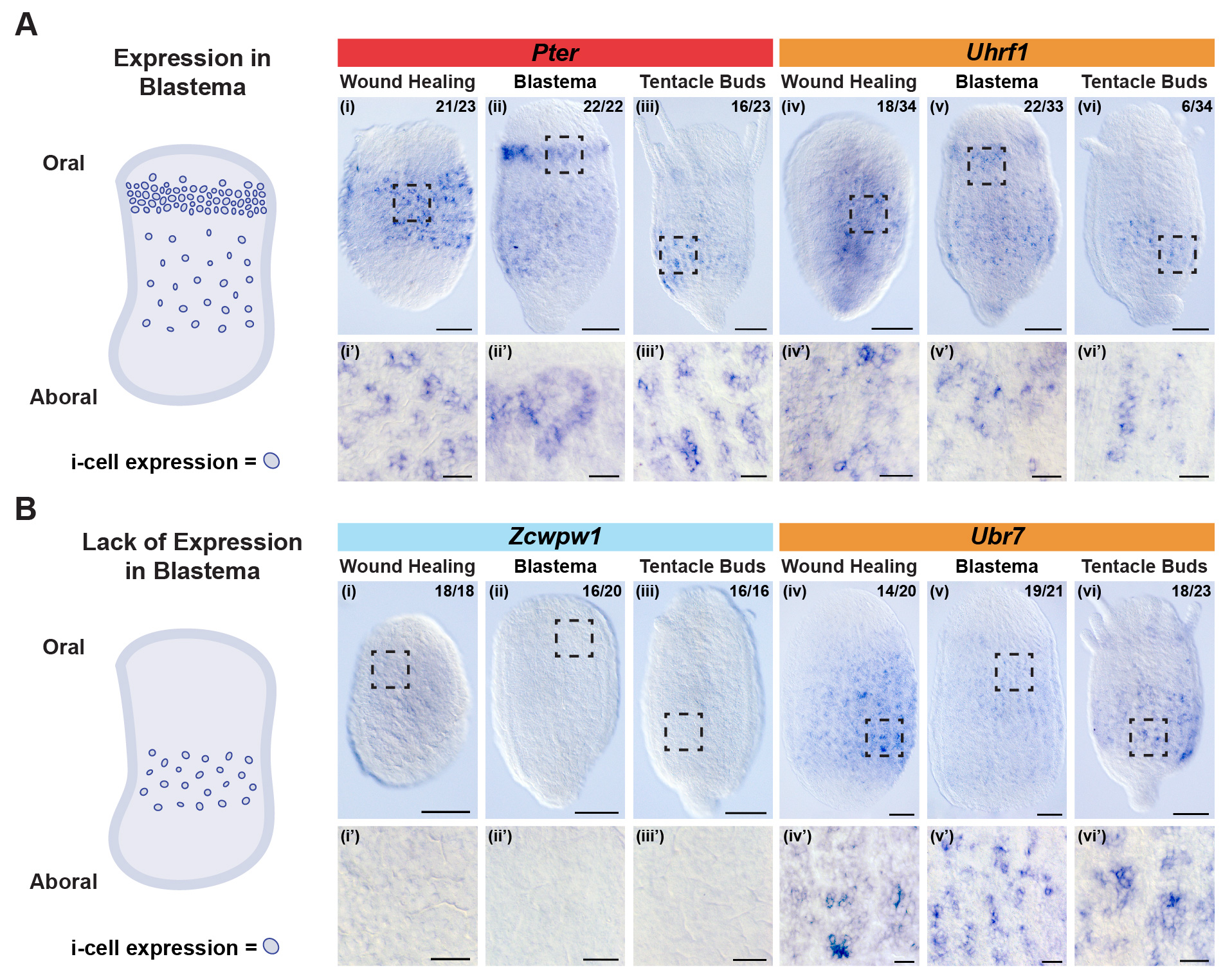

### S4 Fig

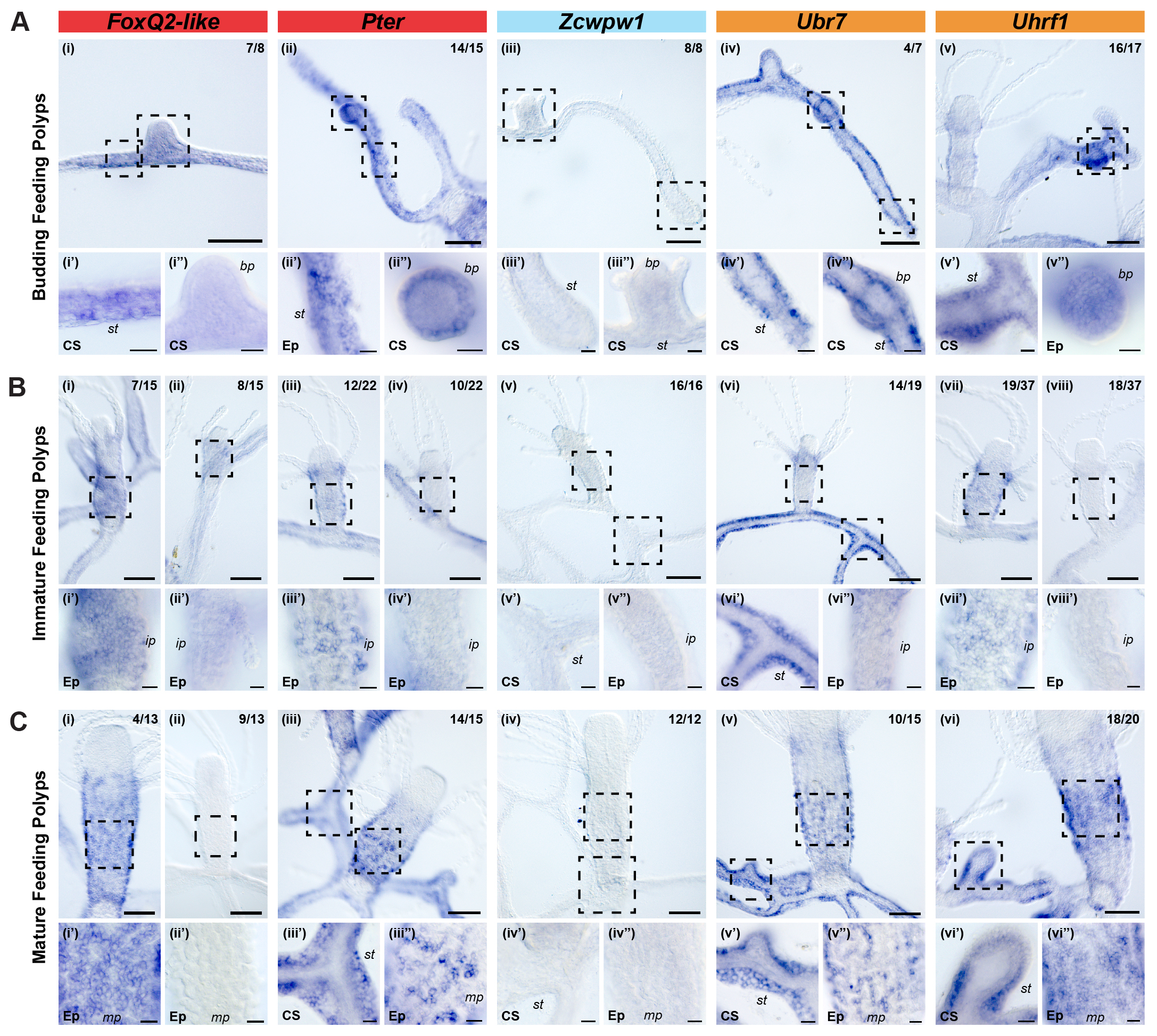

### S5 Fig

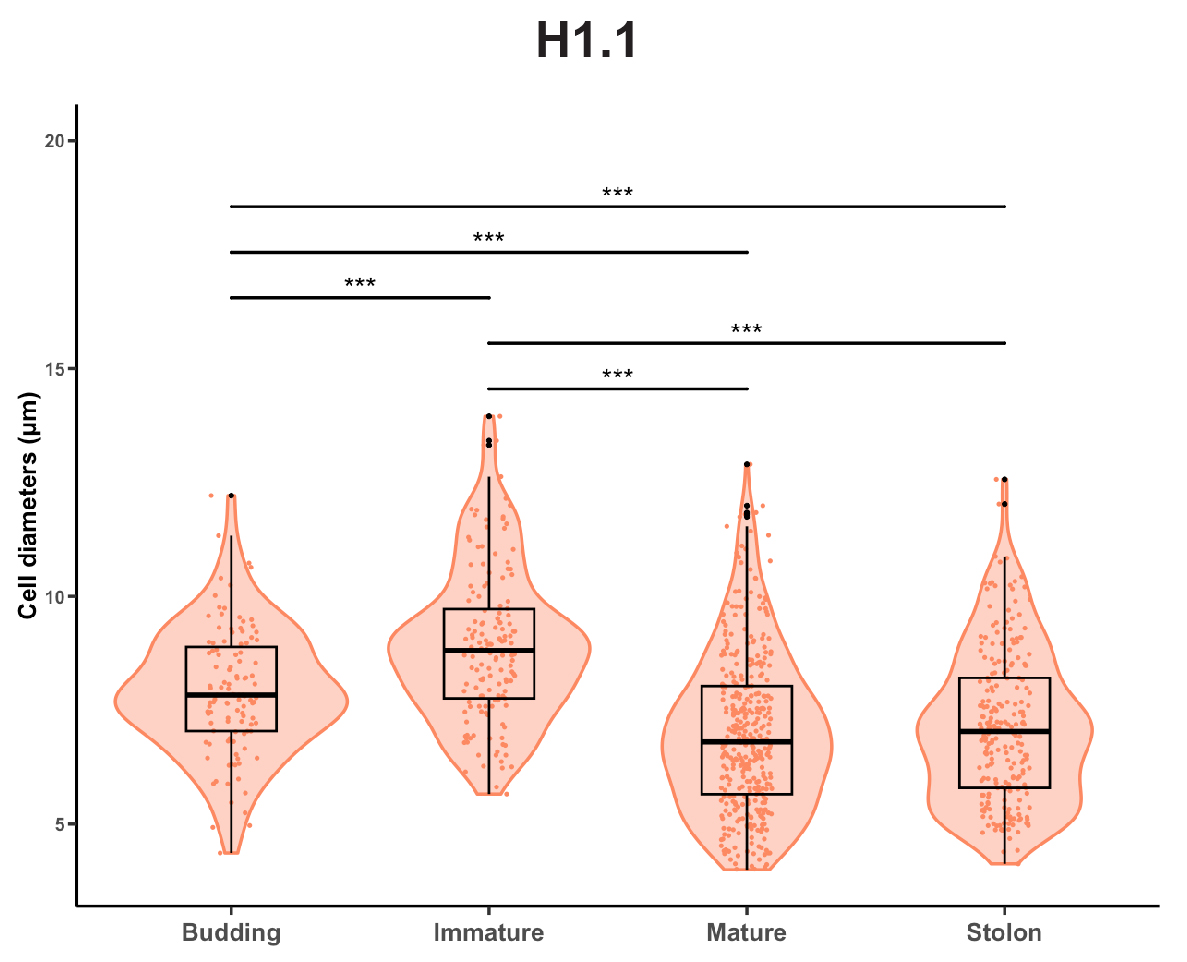

### S6 Fig

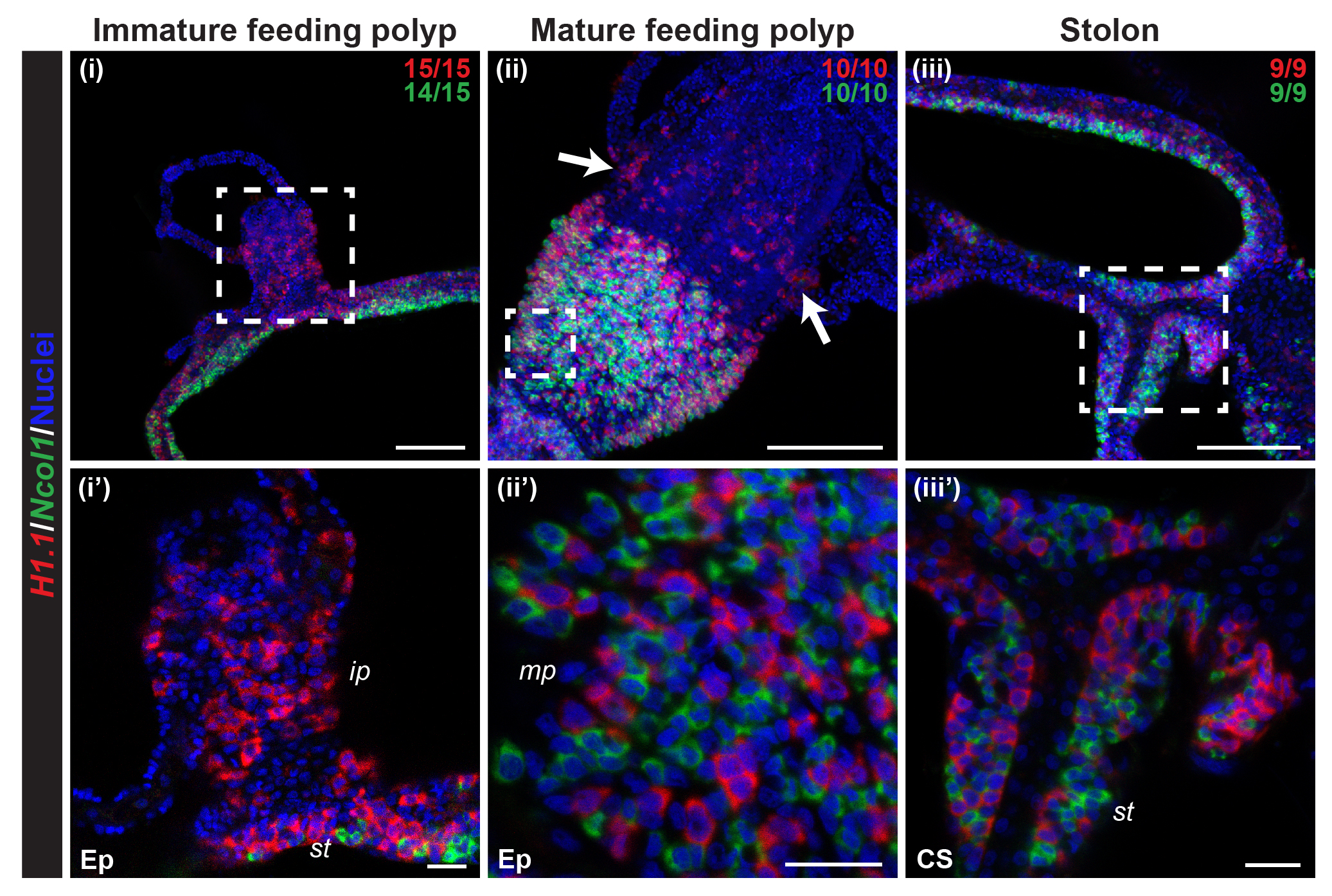
